# The genetic origin of evolidine, the first cyclopeptide discovered in plants, and related orbitides

**DOI:** 10.1101/2020.06.10.145326

**Authors:** Mark F. Fisher, Colton Payne, Thaveshini Chetty, Darren Crayn, Oliver Berkowitz, James Whelan, K. Johan Rosengren, Joshua S. Mylne

**Affiliations:** The University of Western Australia, School of Molecular Sciences & The ARC Centre of Excellence in Plant Energy Biology, 35 Stirling Highway, Crawley 6009, WA, Australia; The University of Queensland, Faculty of Medicine, School of Biomedical Sciences, St Lucia, Queensland 4069, Australia; Australian Tropical Herbarium, James Cook University, PO Box 6811, Cairns QLD 4870, Australia; Department of Animal, Plant and Soil Sciences, School of Life Sciences & ARC Centre of Excellence in Plant Energy Biology, AgriBio, The Centre for AgriBioscience, La Trobe University, Bundoora 3086, VIC, Australia

**Author notes:** Corresponding author: Joshua S. Mylne.

## Abstract

Cyclic peptides are reported to have antibacterial, antifungal and other bioactivities. Several genera of the Rutaceae family are known to produce orbitides, which are small head-to-tail cyclic peptides composed of proteinogenic amino acids and lacking disulfide bonds. *Melicope xanthoxyloides* is an Australian rain forest tree of the Rutaceae family in which evolidine - the first plant cyclic peptide - was discovered. Evolidine (cyclo-SFLPVNL) has subsequently been all but forgotten in the academic literature, but here we use tandem mass spectrometry to rediscover evolidine and using *de novo* transcriptomics we show its biosynthetic origin to be from a short precursor just 48 residues in length. In all, seven *M. xanthoxyloides* orbitides were found and they had atypically diverse C-termini consisting of residues not recognized by either of the known proteases plants use to macrocyclize peptides. Two of the novel orbitides were studied by nuclear magnetic resonance spectroscopy and although one had definable structure, the other did not. By mining RNA-seq and whole genome sequencing data from other species, it was apparent that a large and diverse family of peptides is encoded by sequences like these across the Rutaceae.

## Introduction

Orbitides are a group of cyclic peptides found in plants. They contain between five and 16 amino acid residues, are homodetic cycles and have no disulfide bonds (1,2). Orbitides are thought to be ribosomally-synthesized and post-translationally modified peptides (RiPPs). Their genetic origins are known only for a few groups: the segetalins of *Vaccaria hispanica* (3), some linusorbs (previously cyclolinopeptides) from flax (*Linum usitatissimum*) (4), two curcacyclines from *Jatropha curcas* (1), a number of orbitides in *Citrus* species (3,5), the annomuricatins of *Annona muricata* (6) and the PawL-derived peptides (PLPs) from members of the daisy family (Asteraceae) (7–9).

Evolidine is an orbitide from the leaves of *Melicope xanthoxyloides* (F. Muell.) T.G. Hartley (synonym *Euodia xanthoxyloides* F. Muell.), a species of the Rutaceae family which is found in the rainforest of northern Queensland, Australia, the Lesser Sunda Islands, the Moluccas, Timor Leste, and New Guinea (10–12). Evolidine was first isolated at the University of Sydney, Australia in 1952 (13) and its amino acid content and cyclic nature were determined in 1955 (14). The amino acid sequence was confirmed in 1961 (15) and the first synthesis of the peptide was carried out in 1965 (16). A number of subsequent studies of the structure of evolidine have been carried out, using both NMR (17,18) and X-ray crystallography (**Fig. 1**) (19). In 2005, evolidine was reported to have antibacterial and antifungal activity (20).

**Fig. 1.**
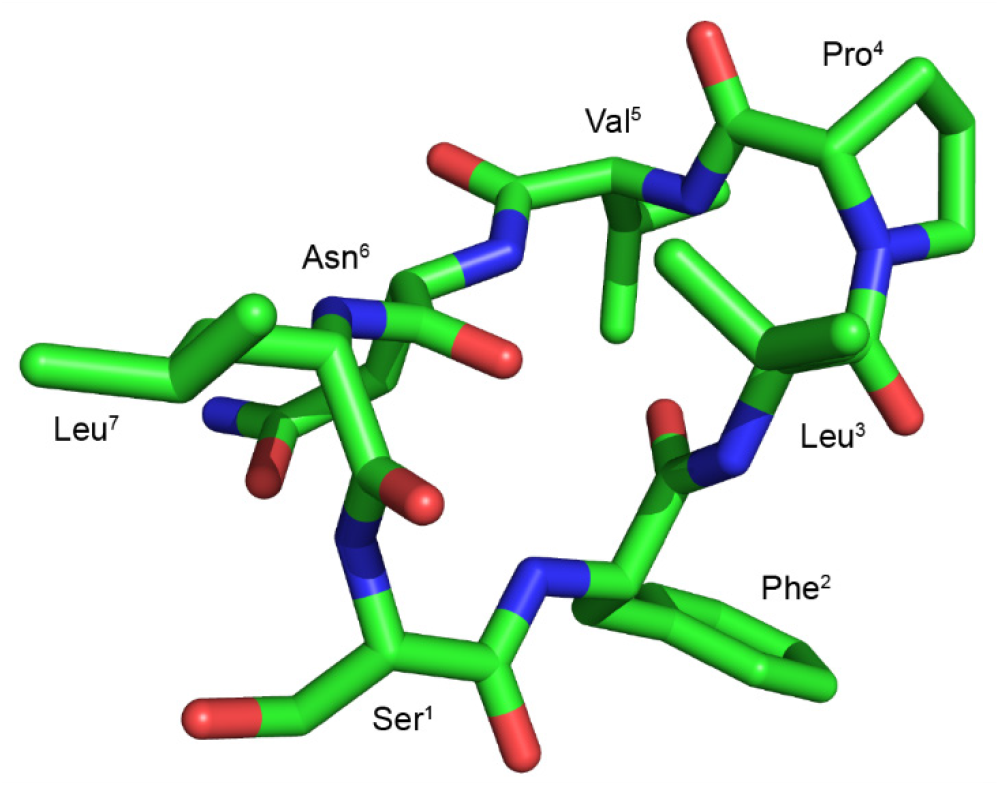
Structure of evolidine. Deposition no. 1266027 at the Cambridge Structural Database, obtained by X-ray crystallography (19).

Despite being the first cyclic peptide isolated from a plant, evolidine has been overlooked by the natural product literature. It was not included in the most comprehensive review of plant cyclic peptides to date (21). Instead - perhaps due to the economic importance of flaxseed, from which it comes - cyclolinopeptide A is often cited as the first cyclic peptide (or sometimes the first orbitide) discovered in a plant (1,4,21–25). However, cyclolinopeptide A was not isolated until 1959 (26) and its chemical structure solved only in 1966 (27).

We were interested in investigating the biosynthesis of evolidine, the first plant cyclopeptide, and so we assembled a *de novo* transcriptome for leaves of *M. xanthoxyloides*. We found mRNA transcripts that encode evolidine and used mass spectrometry to rediscover evolidine plus confirm six novel orbitides encoded by similar transcripts. The C-terminus of these core peptides were Phe, Lys, Leu and Ser - none of which are resides recognized by either of the known plant macrocyclizing enzymes; namely asparaginyl endopeptidase of prolyl oligopeptidase. Two of the novel orbitides were examined by NMR spectroscopy and their behavior varied in solution. Using the precursor sequences from *M. xanthoxyloides*, we analyzed RNA-seq and whole genome sequencing (WGS) data from the Sequence Read Archive (28) (SRA) and found genes and transcripts encoding putative peptides in other Rutaceae species.

## Results and Discussion

We first confirmed the presence of evolidine in our tissue sample by liquid chromatography-mass spectrometry (LC-MS). The peptide was sequenced by tandem mass spectrometry (MS/MS) (**Fig. S1**). To find the transcript that encodes evolidine, we extracted total RNA from *M. xanthoxyloides* leaves, performed RNA-seq and assembled a transcriptome using established methods (7). We searched the transcriptome for sequences encoding evolidine by using cyclic peptide precursor sequences from *Citrus* species; we reasoned that these could share sequence similarity with the putative gene coding for evolidine because both *Melicope* and *Citrus* belong to the Rutaceae family. We found two sequences potentially coding for evolidine along with similar, putative peptide-encoding sequences (**Fig. 2**). Aligning these sequences showed absolute conservation for the residues flanking evolidine (Glu and Ser), which indicated where the peptide sequence might lie. We named the potential cyclic peptides xanthoxycyclins and the genes *Proxanthoxycyclins*.

**Fig. 2.**
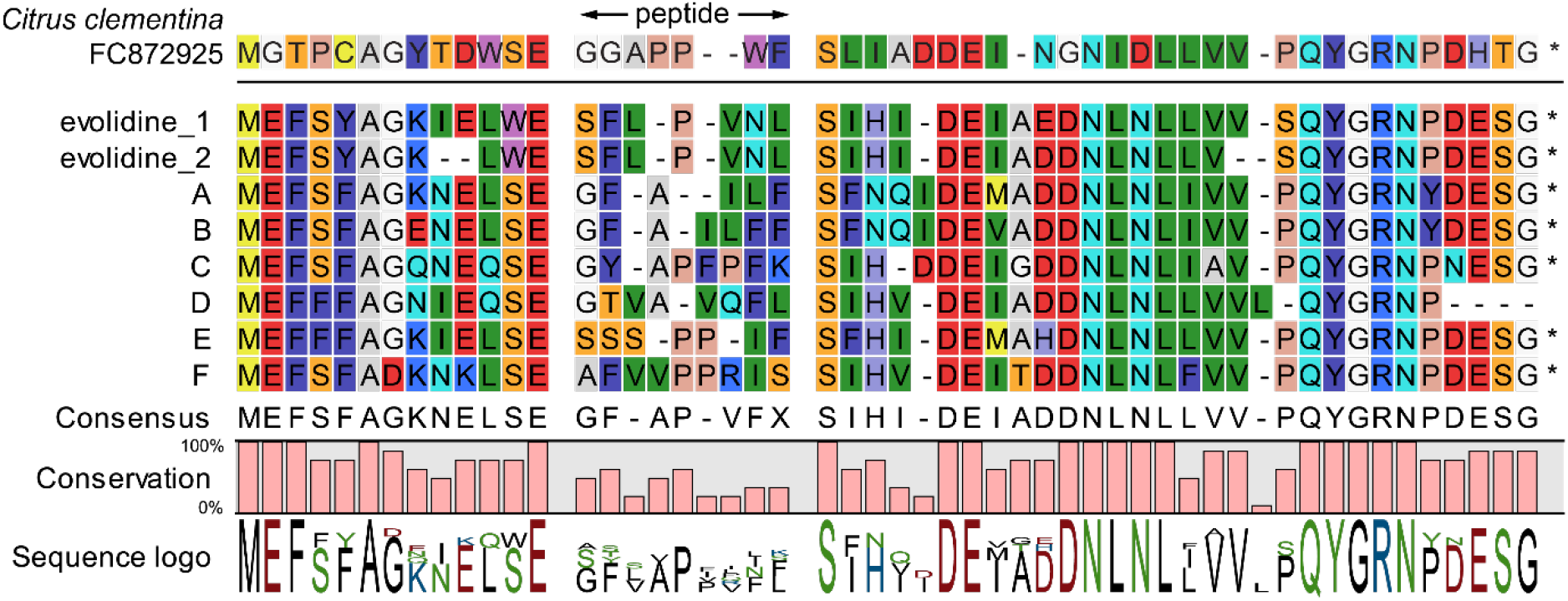
*M. xanthoxyloides* peptide precursors aligned with a known *Citrus* orbitide precursor. The translated coding sequences for seven cyclic peptides in *M. xanthoxyloides* aligned to *Citrus clementina* Genbank accession FC872925. The letters A to F stand for *Proxanthoxycyclins A* to *F*.

We confirmed the sequences obtained by RNA-seq and transcriptome assembly by designing primers against the ends of each contig to amplify either genomic DNA, or a full length coding sequence using cDNA produced from poly-T-primed RNA as the template. We were able to confirm one of the *Proevolidine* sequences (**Fig. 3**), and *Proxanthoxycyclin E* (**Fig. S2A**) from genomic DNA. *Proxanthoxycyclins B-D* were sequenced from cDNA and found to be a 100% match to the contigs assembled by RNA-seq. It was not possible to Sanger sequence *Proxanthoxycyclin A* due to the coding sequence being almost identical to that of *Proxanthoxycyclin B* and the repetitive nature of one of the regions to be primed. We were also unable to PCR amplify the putative second *Proevolidine* gene or clone a sequence for *Proxanthoxycyclin F*.

**Fig. 3.**
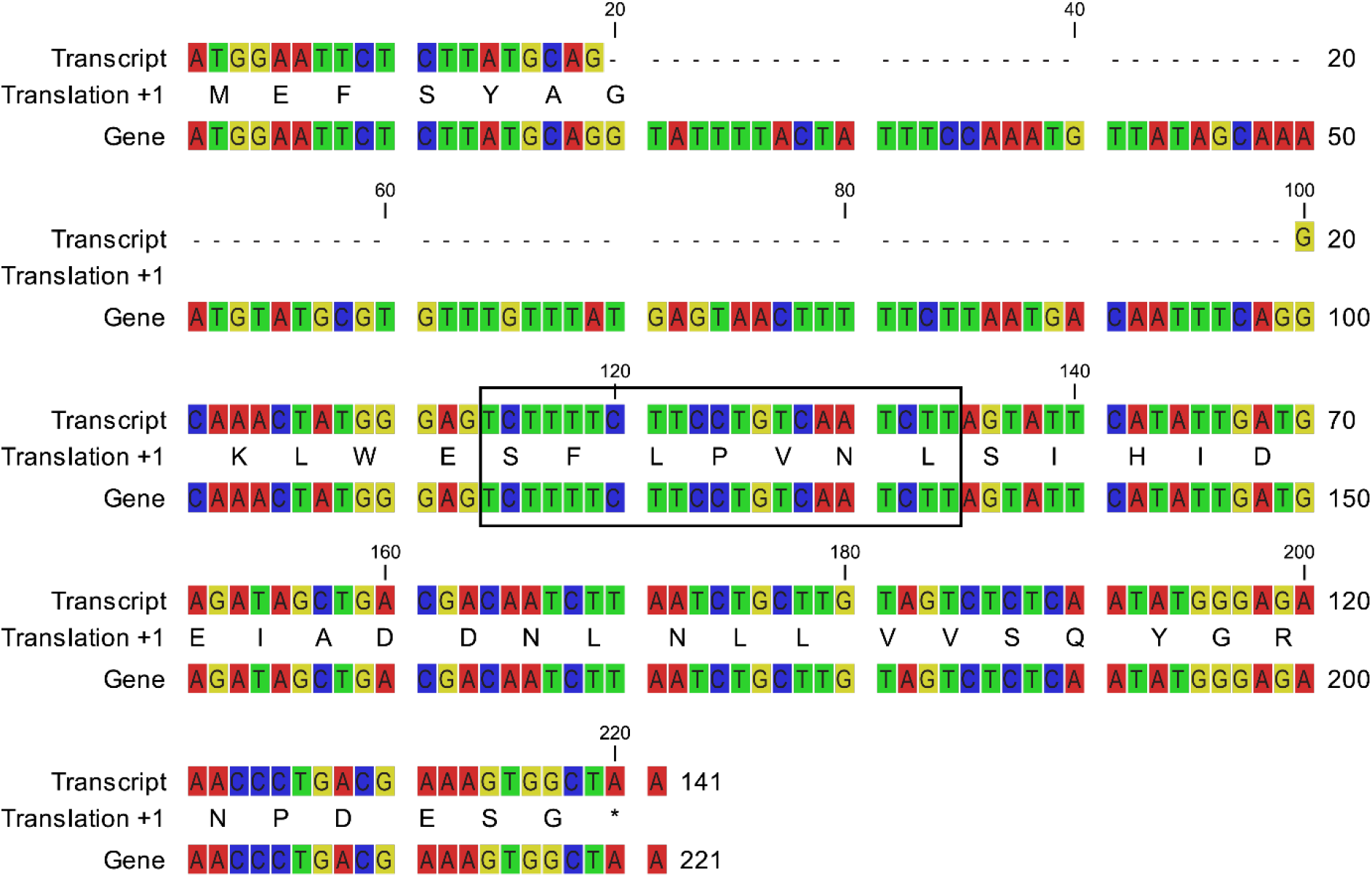
Alignment of *Proevolidine* transcript and gene. The Sanger-sequenced gene data matches perfectly with the RNA-seq transcript, except that the gene contains an 80 bp intron. The translated peptide sequence is shown between the nucleotide sequences; evolidine is shown within a box.

### Confirmation of further cyclic peptides by tandem mass spectrometry

Having the sequences for transcripts encoding novel cyclic peptides facilitated their identification and sequencing from LC-MS/MS data. We found peaks corresponding to the six additional peptides predicted by transcriptomic data and sequenced the peptides by MS/MS (**Table 1**). We named these peptides xanthoxycyclins A to F. The LC-MS data for xanthoxycyclin A are shown in **Fig. 4**; data for xanthoxycyclins B to F are shown in **Fig. S3–S7**.

**Table 1.**
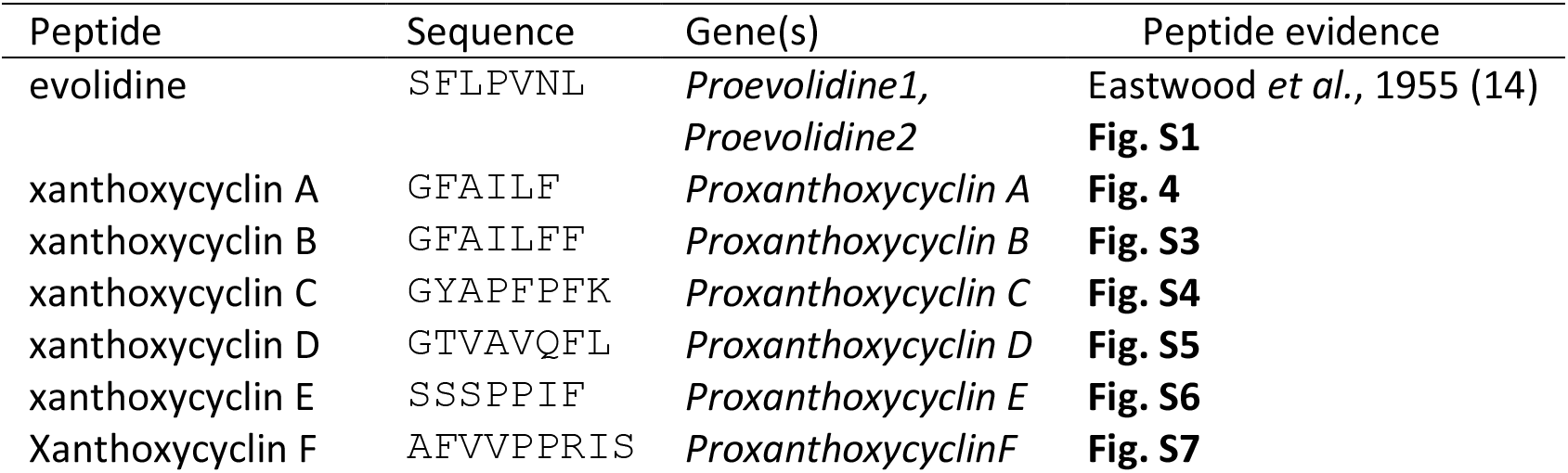
Summary of *M. xanthoxyloides* cyclic peptides, the genes that encode each and evidence for the peptides from mass spectrometry. The peptides are all backbone cyclic, with the sequences beginning with their proto-N-terminus and ending with their proto-C-terminus that are joined during processing.

**Fig. 4.**
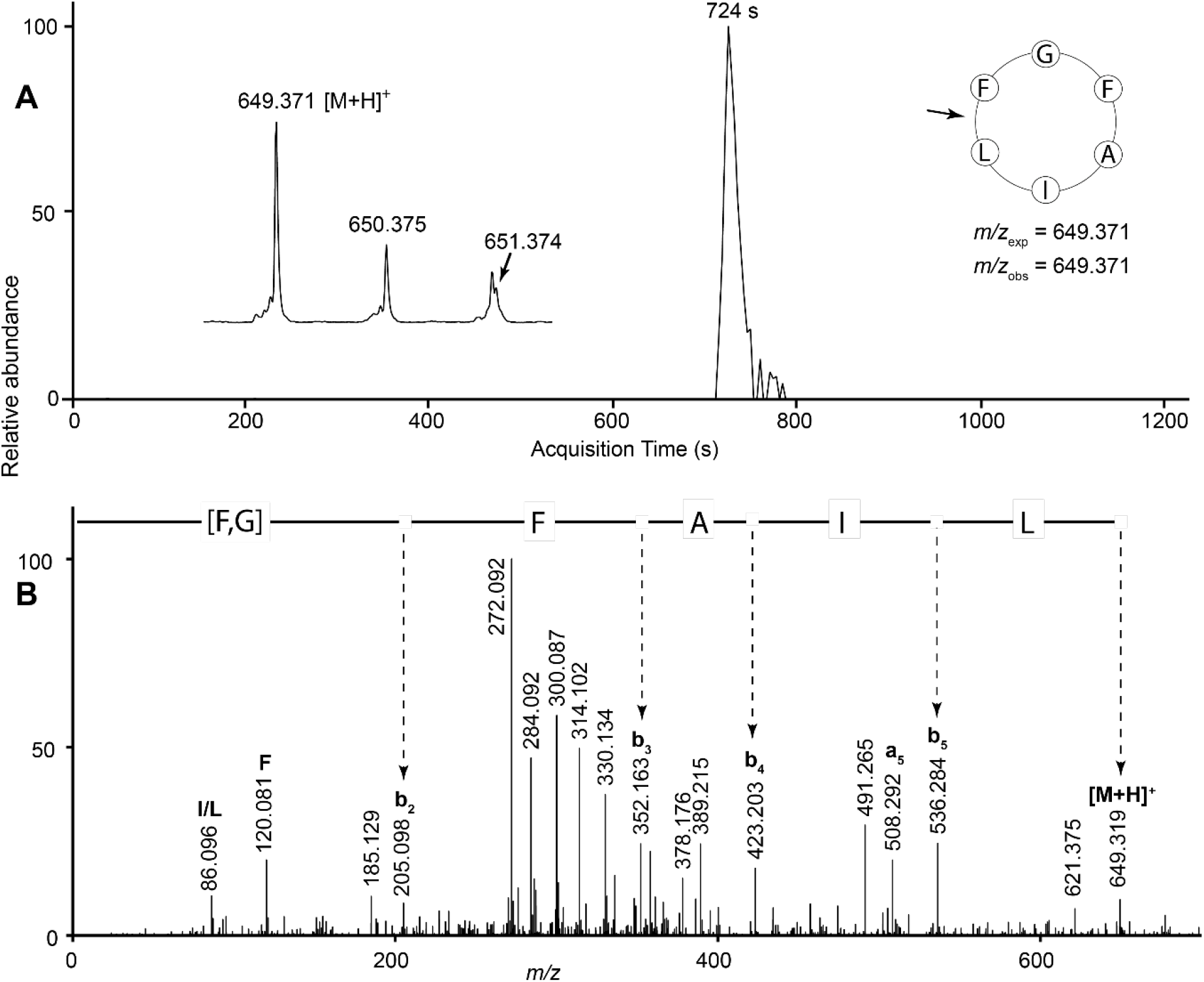
LC-MS/MS data for xanthoxycyclin A. (A) Extracted ion chromatogram at *m/z* 649.371. Inset left: mass spectrum at the chromatogram peak. Inset right: cyclic peptide sequence with MS/MS ring cleavage point shown by an arrow, expected *m/z* (*m/z*_exp_) and observed *m/z* (*m/z*_obs_). (B) MS/MS spectrum of protonated molecule at *m/z* 649.371. A strong sequence of b ions is derived from initial ring opening between Leu and Phe residues. Immonium ions are labelled with the corresponding amino acid residue.

The sequences of the core peptides lack conservation except some at the N and C-terminal residues: the N-terminal residue were ones with small side chains, namely Gly, Ser and Ala. The C-terminal residue was one of four possibilities; either Phe or Leu, Lys for xanthoxycyclin C and Ser for xanthoxycyclin F. In the sequence preceding the peptides, Glu is absolutely conserved in the P1 position and the P2 residue is a highly-conserved Ser, although for the two evolidine transcripts a Trp is encoded. There are many other highly conserved residues, especially in the sequence following the core peptide, where the P1’ residue is always Ser and P2’ is most often Ile (**Fig. 2**). The core peptides described above are typical orbitides in that they contain mainly hydrophobic residues (1). With between six and eight residues, they fall into the low-middle range of sizes for orbitides, which can be as large as 16 amino acids (2).

It is perhaps worth noting that when amplifying *Proxanthoxycyclin E* from genomic DNA template, as well as finding the sequence we were searching for, we also amplified six other sequences using the same primers (**Fig S2B**). Five of these were complete ORFs with a high degree of sequence similarity to the *Proxanthoxycyclins;* another appeared to be a truncated form. There was no evidence to support these in the leaf MS/MS data, but these genes related to *Proxanthoxycyclin* might not be expressed or could encode xanthoxycyclins that are expressed in other tissues or specific growth stages not represented within the leaf sample we analyzed.

### Potential peptide-encoding transcripts and genes in other Rutaceae species

To investigate the prevalence of this kind of peptide-encoding transcript within the Rutaceae, we searched the SRA to find RNA-seq and WGS data for other species in the family. We downloaded and assembled RNA-seq data from *Aegle marmelos* fruit pulp, *Atalantia buxifolia* mature fruit, *Bergera koenigii* leaf, *Clausena excavata* leaf*, Zanthoxylum armatum* young leaf and *Zanthoxylum bungeanum* young leaf, as well as WGS data for *Atalantia buxifolia* and *Clausena lansium*. No peptide-encoding transcripts were detected in either *Zanthoxylum* species despite their being closely related to *Melicope* and there being three known orbitides in three different species in the genus *Zanthoxylum* (29–31). Putative peptide-encoding transcripts were found in all other RNA-seq datasets examined (**Fig. 5**).

**Fig. 5.**
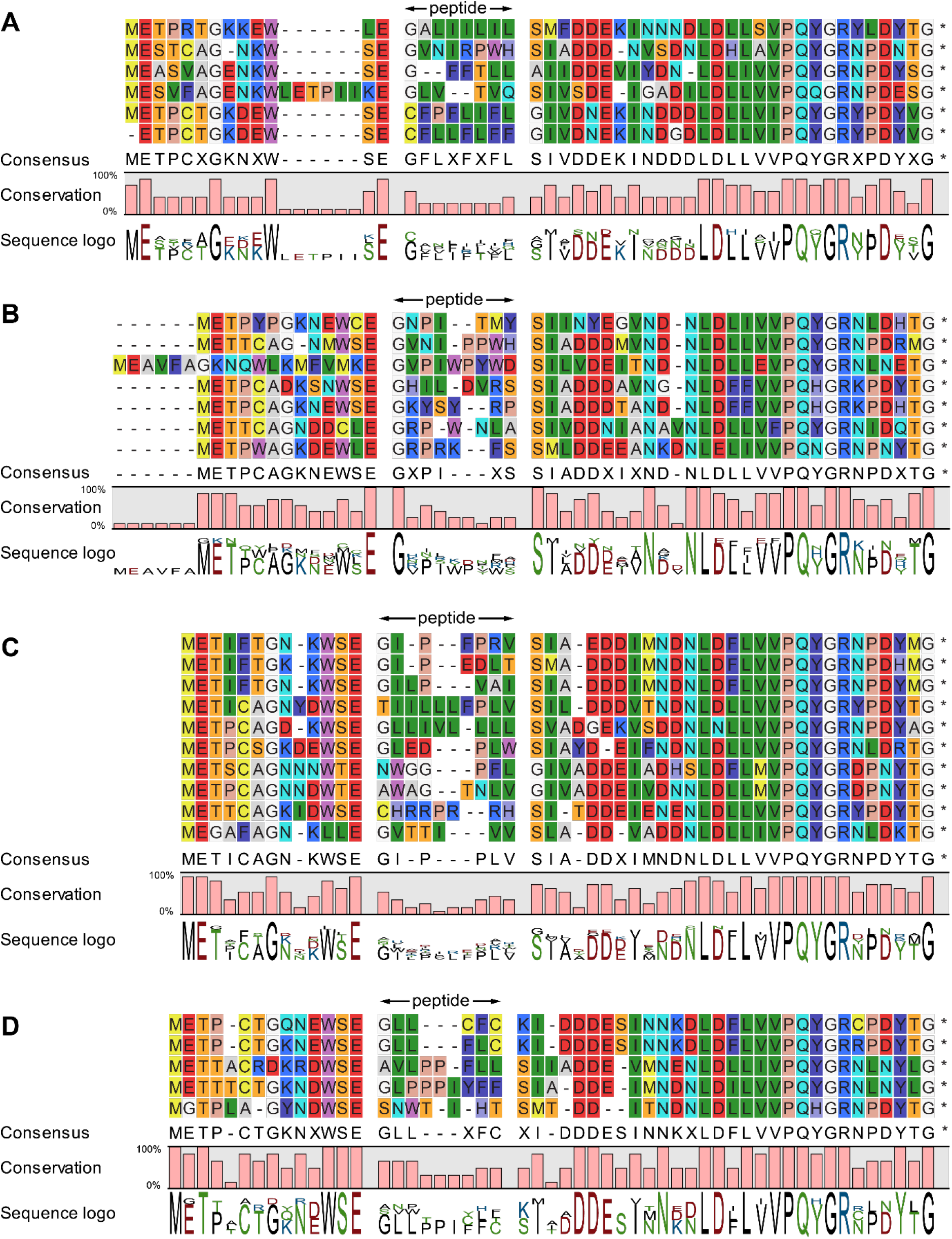
Alignments of putative propeptide sequences from four Rutaceae species. Sequences are from (A) *Aegle marmelos;* (B) *Atalantia buxifolia*, (C) *Bergera koenigii* and (D) *Clausena excavata*. Sequences are shown with background RasMol colours, and the putative peptide is marked and spaced from the surrounding sequence. Asterisks represent stop codons. Below each alignment is the consensus sequence, the degree of conservation and a sequence logo.

Like those of *M. xanthoxyloides*, the potential propeptides encoded within these four species also show a high sequence similarity outside the core peptide sequence. Within the core peptide, Gly is the most common residue at the N-terminus, but the C-terminual residues are even less conserved than the core peptides of *M. xanthoxyloides*. In *A. marmelos*, Phe and Leu are most common, as in *M. xanthoxyloides*, but the other three species have a wide range of C-terminal residues, including Val, Thr, Trp, Pro and Ser, among others. It must be noted, however, that there is no peptide evidence yet to indicate that these encoded sequences are actually processed into orbitides, so it would be unwise to speculate overly. The high degree of conservation, with up to 50% identity and 60% similarity between the flanking regions of the *M. xanthoxyloides* propeptides and the *Citrus clementina* sequence FC872925, suggests that these regions are important for peptide processing.

WGS data from the species *A. buxifolia* and *C. lansium* contained many potential peptide-encoding sequences (**Fig. S8** and **Fig. S9** respectively), though it was not possible to say how many might be expressed without transcriptomic data. One of the sequences from *C. lansium* appeared to encode the known peptide clausenlanin A (cyclo-GLILLLLLL) previously found in that species (32). Another sequence potentially encoding a peptide from *C. lansium*, clausenlanin B (cyclo-GLVLLLLLL) (32), was found, but not in the data for that species; rather, it was found in WGS data from *A. buxifolia*. The two putative propeptide sequences show a remarkably high degree of conservation, differing only at three residues outside the core peptide sequence (**Fig. S10**). The strong degree of sequence similarity in the propeptides shown among the species studied here and *Citrus* species investigated elsewhere (3,5), especially in the sequence following the core peptide, suggests a common point of evolution for all these peptides across the Rutaceae.

### NMR spectroscopy of synthetic xanthoxycyclins

Orbitides, due to their small size and restrained nature, contain a limited variety of structural features. The majority of orbitides are five to ten residues with structural features being limited to ordered loops (2), turns (33,34) and mini-β-sheets (35). Despite over 100 orbitides being discovered and reported in CyBase (36), less than ten 3D structures have been solved using either solution NMR spectroscopy or X-ray crystallography and submitted to the Protein Data Bank (PDB) (36).

In this work we wanted to further explore the structural features of orbitides and so we subjected synthetic xanthoxycyclin D and xanthoxycyclin F to solution NMR spectroscopy. The 1D NMR spectrum of xanthoxycyclin D demonstrated sharp peaks with good signal dispersion in H2O/D2O. A set of 2D NMR spectra, including TOCSY (Total Correlation Spectroscopy) (37) and NOESY (Nuclear Overhauser Effect Spectroscopy) (38), were recorded and assigned using a sequential assignment strategy. The NOESY spectrum contained a large number of medium range NOEs (**Table 2**), consistent with the peptide adopting an ordered structure. Xanthoxycyclin D demonstrated relatively small deviations from random coil Hα shifts (**Figure S11**), however, deviations in Cα and Cβ chemical shifts (**Figure S11**) allowed for prediction of backbone dihedral angles by TALOS-N (Torsion Angle Likelihood Obtained from Shift and sequence similarity) (39) for seven of the eight residues.

**Table 2.** NMR structure statistics of xanthoxycyclin D

|  |  |
| --- | --- |
| **Energies (kcal/mol)** |  |
| Overall | -225.70 ± 0.18 |
| Bonds | 1.88 ± 0.01 |
| Angles | 5.43 ± 0.06 |
| Improper | 2.16 ± 0.03 |
| Dihedral | 34.76 ± 0.03 |
| Van der Waals | -16.82 ± 0.08 |
| Electrostatic | -253.16 ± 0.19 |
| NOE | 0.006 ± 0.001 |
| Cdih | 0.036 ± 0.002 |
| **MolProbity statistics** |  |
| Clashes (>0.4 Å / 1000 atoms) | 0 |
| Poor rotamers | 0 |
| Ramachandran outliers (%) | 0 |
| Ramachandran favored (%) | 100 ± 0 |
| MolProbity score | 0.5 ± 0 |
| MolProbity score percentile | 100 ± 0 |
| **Atomic RMSD (Å)** |  |
| Mean global backbone | 0.20 ± 0.06 |
| Mean global heavy | 0.93 ± 0.24 |
| **Experimental restraints** |  |
| **Distance restraints** |  |
| Short range (i-j < 2) | 50 |
| Medium range (/i-j/ < 5) | 24 |
| Long range (/i-j/ > 5) | 0 |
| Hydrogen bond restraints | 8 (4 H-bonds) |
| Total | 82 |
| **Dihedral angle restraints** |  |
| φ | 7 |
| ψ | 7 |
| χ<sup>1</sup> | 1 |
| Total | 15 |
| **Violations from experimental restraints** |  |
| Total NOE violations exceeding 0.2 Å | 0 |
| Total dihedral violations exceeding 2.0° | 0 |

The dihedral angle restraints and NOE based distance restraints were used as input for simulated annealing and energy minimization in explicit water, and resulted in a tightly overlapping structural ensemble with an RMSD of 0.20 Å for the backbone atoms and 0.93 Å for all heavy atoms (**Fig. 6**). Only one residue, Gln6, appears to have complete conformational freedom of its sidechain. The sidechain of Phe7 adopts two conformational states; these states are reflected in the NMR data, which show a large chemical shift separation of the two HB protons but averaged ^3^J_HαHβ_ coupling constants. The sidechain of Leu8 is locked into a singular conformation (gauche-trans); again reflected in the NMR data, which shows a slightly weaker NH-Hβ2 NOE compared to the NH-Hβ3 NOE and a large difference in ^3^J_HαHβ_ coupling constants, with the coupling between Hα and Hβ2 being significantly weaker than that between Hα and Hβ3 (40).

**Fig. 6.**
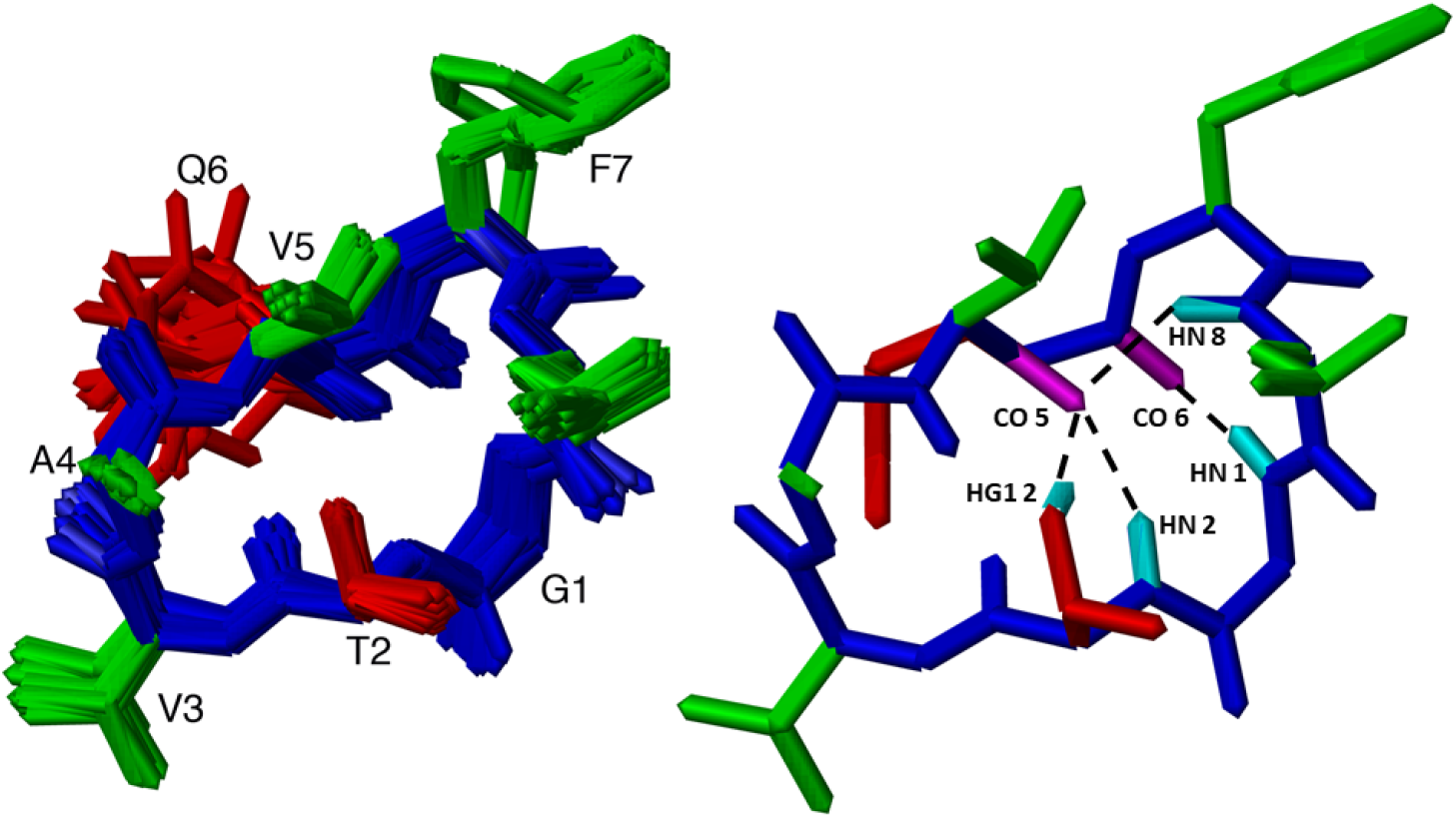
NMR solution structure of xanthoxycyclin D. Left image is of the structural ensemble of 20 best structures superimposed, shown in stick format. Right image is a single structure of xanthoxycyclin D in stick format illustrating hydrogen bonding with dashed lines. Hydrogen bond donors are shown in cyan, with hydrogen bond acceptors being shown in magenta. Both left and right images show the peptide backbone in blue, hydrophobic sidechains in green and polar sidechains in red.

The structure of xanthoxycyclin D has two notable features: First, a typical type III β-turn between residues Val5 and Leu8. The Type III turn is defined by a distance between the Cα carbons of Val5 and Leu8 (*i* and *i*+3) of 4.5 Å, with the ϕ and ψ dihedral angles of Gln6, −62°, −34° (*i*+1), and Phe7, −60°, −31° (*i*+2), respectively. The turn is stabilized by a hydrogen bond between the NH of Leu8 and CO of Val5. Second, an atypical type III β-turn is formed between Thr2 and Val5. Although the distance between the Cα carbons of Thr2 and Val5, which is 5 Å, is consistent with a turn, and the ϕ and ψ dihedral angles of Val3, −79°, −31° (*i*+1) and Ala4, −79°, −39° (*i*+2) are similar to the common type III β-turn angles, the hydrogen bond which stabilizes this turn is formed between the HN of Val5 and the side chain OG1 of Thr2 rather than the backbone carbonyl. Hydrogen bonds are also present between Gly1 HN - Gln6 O and Thr2 HN – Val5 O. Additionally, the hydroxyl proton HG1 of Thr2 is within hydrogen bond distance of the carbonyl of Val5 in half of the models of the structural ensemble. A weak resonance at 5.15 ppm was indeed detected and assigned to Thr2 HG1 in the NMR data, which is consistent with it being involved in a hydrogen bond. However, this implies that there are three protons potentially competing to form hydrogen bonds with the same carbonyl oxygen, thus rearrangements allowing this sharing of acceptor would likely be required.

Xanthoxycyclin F was also studied using solution NMR spectroscopy. In contrast to xanthoxycyclin D, the 1D spectrum was of poor quality, and despite the appearance of a single pure compound on HPLC analysis, the peptide adopted multiple conformations in solution, evident from the numerous NMR peaks observed. The 2D TOCSY and NOESY data confirmed numerous conformations at almost every residue, presenting a challenge for the assignment of the data. Despite this, a dominant conformation was able to be assigned and confirmed using a natural abundance ^13^C HSQC (Heteronuclear Single Quantum Coherence spectroscopy). The secondary shifts of the dominant confirmation were determined for all residues of xanthoxycyclin F (**Figure S12**). It demonstrated secondary shifts that deviate from random coil for most residues, but the complete lack of any non-sequential NOEs prevented the determination of a structure for any of the conformers. A previous study showed that some members of the PLP subfamily of orbitides (PLPs −2, −4, −10 & −12) also demonstrate secondary Hα shifts which deviate from random coil (8), indicating at least some degree of ordered backbone, yet they displayed only irregular non-rigid structures lacking classical secondary structural features.

### Biological activities of *M. xanthoxyloides* orbitides

Poojary *et al*. (20) reported that evolidine inhibits the growth of some bacteria and fungi. We attempted to replicate these results and to extend the study to xanthoxycyclins D and F, using disc diffusion assays like those in the prior work. Our synthetic peptides were generated in house by solid phase peptide synthesis or purchased so that we had enough material for NMR structure determination and to test the antifungal and antibacterial activity of the peptides. LC-MS/MS was used to compare the properties of the natural and synthetic compounds to verify that they were identical. The LC retention times were compared and mirror plots of the MS/MS spectra for each natural/synthetic pair were made. We concluded that the natural and synthetic compounds were identical. This was done for all three synthetic peptides: evolidine (**Fig. S13**), xanthoxycyclin D (**Fig. S14**) and xanthoxycyclin F (**Fig. S15**).

Our results showed no growth inhibition by any of the three peptides against the Gram-negative bacterium *Escherichia coli* K-12 nor against the Gram-positive bacterium *Bacillus subtilis* Marburg No. 168, whereas a positive control disc containing 50 μg of kanamycin showed a clear zone of inhibition in every case (**Fig. S16**). Similarly, the peptides were not active against either *Candida albicans* or *Aspergillus fumigatus* (**Fig. S17**) although a positive control disc containing 50 μg of amphotericin B inhibited fungal growth. We attempted to use griseofulvin as a positive control to follow the method of Poojary *et al*., but found it had no effect on either fungal species, even with a quantity four times that used in the earlier study (**Fig. S18**).

Our results stand in contrast to the prior work (20), which found that evolidine inhibited growth of *B. subtilis* (strain not specified), and the fungi *C. albicans* and *Aspergillus niger* using a similar assay to ours. The same study was consistent with our finding that *E. coli* is unaffected by evolidine. The prior study did not state what quantities of evolidine were applied to the discs, preventing a direct comparison of the results. This is not the first example of cyclic peptide bioactivity being non-reproducible, though this occurred mostly when synthetic compounds were compared to natural products. This was previously observed in yunnanins A and C (orbitides from *Stellaria yunnanensis*), where the synthetic compounds had activity orders of magnitude lower than the natural products (41). The authors attributed this either to conformational differences between the natural and synthetic products, or traces of another compound in the natural extract. A similar phenomenon has been seen for other, non-orbitide cyclic peptides, such as phakellistatins 1 and 10 (41), phakellistatin 11 (42) and dolastatin 16 (43). This last case is intriguing, because the study compared the synthetic product with natural samples extracted and tested previously (44). The natural samples were repurified using the same method applied to the synthetic compound and their activity was found to be greatly reduced compared to the earlier results (43,44). This lends weight to the hypothesis that trace impurities may be responsible for bioactivities found only in natural extracts.

In our case, the compounds in our study and that of Poojary *et al*. were both synthetic, but differences in the method of purification could account for the observed differences in activity. Poojary *et al*. (20) purified evolidine using silica gel chromatography followed by recrystallization from EtOAc-*n*-hexane, whereas we performed two rounds of reversed phase-high performance liquid chromatography (RP-HPLC).

### Biosynthesis of *M. xanthoxyloides* orbitides

Several studies have investigated the post-translational processing required to mature plant cyclic peptides. Asparaginyl endopeptidase, a protease that recognizes Asp or Asn at the C-terminus of the core peptide for cleavage, has also been recruited by evolution multiple times to perform cyclization of different cyclic peptide families via a transpeptidation reaction. These include (i) kalata-type cyclic peptides (cyclotides), a family of plant macrocyclic peptides containing three disulfide bonds (45–47); (ii) the cyclic knottins – a family of mostly trypsin-inhibitory peptides from the squash family(46,48) and (iii) the single-disulfide PawS-derived peptides of the daisy family (49) and presumably the closely-related PawL-derived peptides, a group of orbitides in the daisy family with an absolutely conserved Asp at the C-terminus of the core peptide (7,8).

Previous work has revealed that a second cyclization mechanism in plants involving a prolyl oligopeptidase named peptide cyclase 1 (PCY1) is responsible for cyclization of the segetalin orbitides in *Vaccaria hispanica (Saponaria vaccaria*) (3,23). This enzyme recognizes Pro, or sometimes Ala, at the C-terminus of the core peptide. It is likely that a prolyl oligopeptidase is similarly responsible for cyclizing the annomuricatin orbitides in *Annona muricata*, which all have Pro or Ala in an equivalent position (6).

The xanthoxycyclins of *M. xanthoxyloides* differ from both the above groups because their core peptide sequences terminate with Phe, Leu, Ser or Lys. None of these four residues are recognized by the aforementioned asparaginyl endopeptidase (Asn/Asp) or prolyl oligopeptidase (Pro/Ala). If xanthoxycyclins are matured by proteases performing a similar cleavage-coupled intra-molecular transpeptidation with the C-terminal residue, our confirmation of these orbitide sequences indicates one or more novel cyclization mechanisms are yet to be discovered in this species, perhaps involving a different class of protease.

## Conclusion

Here we have shown that evolidine and six novel orbitides, xanthoxycyclins A-F, in the leaves of *M. xanthoxyloides* are genetically encoded and we have solved the solution structure of xanthoxycyclin D by NMR spectroscopy. Using public RNA-seq and WGS data from the SRA, we have also identified similar genes and transcripts in other plants in the Rutaceae family, which may encode comparable cyclic peptides, perhaps from a single evolutionary origin. Our ability to discover all these sequences by searching with peptide precursors from *Citrus* species points to a common ancestor and orbitides made from precursors like these may be present in many of the ∼1,900 species of the Rutaceae (50). A study of other species across this family could reveal more such orbitides that might have interesting pharmacological properties.

## Materials and Methods

### Plant material

Leaves and stems of *M. xanthoxyloides* were collected on 13 June 2019 from Smithfield Conservation Park, Queensland, Australia (16.80609° S, 145.67583° E) under permit PTU18-001474. After removing the stems, the leaves were ground to a powder under liquid nitrogen and stored at −70 °C.

### Synthetic peptides

Evolidine was synthesized using solid phase peptide synthesis on an automated peptide synthesiser (CS Bio Pty Ltd) using fluorenylmethyloxycarbonyl (Fmoc) chemistry on 2-chlorotirityl resin (0.25 mmol scale). Loading of the C-terminal Leu residue (1 M eq.) was achieved by dissolving the FMOC protected Leu residue in dichloromethane, adding four molar equivalents of N,N’-diisopropylethylamine (DIPEA) to the dissolved residue and adding this to the resin. Subsequent amino acids (4 eq.) were activated in 0.5 M 2-(1H-benzotriazole-1-yl)-1,1,3,3-tetramethyluronium hexafluorophosphate and DIPEA (8 eq.) in N,N’-dimethylformamide. The coupling reaction was performed twice for amino acids containing branched β-carbons or aromatic rings. The peptide was cleaved from the resin while maintaining attachment of sidechain protecting groups using 2% trifluoroacetic acid (v/v) in dichloromethane for 3 min. This was repeated ten times. Acetonitrile was added to the peptide solution and dichloromethane and trifluoroacetic acid (TFA) were removed via rotary-evaporation and the peptide portion lyophilized. The peptide was then cyclized in dimethylformamide; this was done by adding 1-[Bis(dimethylamino)methylene]-1H-1,2,3-triazolo[4,5-b]pyridinium 3-oxide hexafluorophosphate (1 eq.) to a 1 mM peptide solution, followed by the slow addition of DIPEA (10 eq.). The reaction was monitored by electrospray ionization mass spectrometry over 3 h. The peptide was then extracted from the dimethylformamide using phase extraction, prior to lyophilization. The remaining sidechain protecting groups were cleaved using TFA/water/triisopropylsilane (97:2:1) for 2 h, the peptide precipitated with cold diethyl ether, filtered and dissolved in 50% acetonitrile before lyophilization. The crude peptide was purified with RP-HPLC on a C18 preparative column (300 Å, 10 μm, 21.20 mm i.d x 250 mm, Phenomenex). Electrospray ionization mass spectrometry was used to confirm the mass of evolidine, with further purifications being conducted on a semi-preparative C18 column (300 Å, 5 μm, 10 mm i.d. x 250 mm, Vydac) with purity being assessed with a C18 analytical column (300 Å, 5 μm, 2.1 mm i.d. x 150 mm, Vydac). Xanthoxycyclin D and xanthoxycyclin F were purchased from GenScript Inc. (Piscataway, NJ) at a purity ≥ 95%.

### Peptide extraction and purification

Peptides were extracted from plants as described previously (7). Briefly, about 1 g of leaf powder was ground with 50 mL of 50% methanol / 50% dichloromethane (v/v). The extract was dried with anhydrous magnesium sulfate and filtered under a vacuum with two 50 mL washes of solvent. After flash chromatography though silica gel, the samples were dried in a rotary evaporator (Heidolph). These leaf peptide extracts were then purified as previously described (8). Briefly, the crude extract was purified by solid-phase extraction using a 30 mg Strata-X polymeric reversed-phase column (Phenomenex). The dried extract was dissolved in 2 mL of an aqueous solution of 5% acetonitrile (v/v) / 0.1% formic acid (v/v) and dispensed onto the column, after which purified peptides were eluted with 75% acetonitrile (v/v) / 0.1% formic acid (v/v). The purified extract was desiccated in a vacuum centrifuge (Labconco) and then prepared for LC-MS analysis by redissolving it in 50 μL of HPLC-grade 5% acetonitrile (v/v) / 0.1% formic acid (v/v) (Honeywell).

### LC-MS/MS for peptide sequencing

Samples (2 μL) were separated by a gradient elution from 5% solvent B to 95% solvent B over 15 min on a high-capacity nano-LC chip (Agilent Technologies; part no. G4240-62010) driven by a 1200 series nano-flow HPLC system (Agilent) at a flow rate of 300 nl/min. Solvent A was 0.1% (v/v) formic acid in water and solvent B was 0.1% (v/v) formic acid in acetonitrile. Both solvents were HPLC-grade (Honeywell). Peptides were detected with either a 6520 or 6550 Q-TOF mass spectrometer (Agilent) via electrospray ionization in positive mode with one MS scan per second. The source voltage was set between 2100 and 2175 V. MS/MS data were gathered by collision-induced dissociation at a rate of two scans per second, with the expected peptide *m/z* values set as “preferred” for fragmentation. Visual inspection of MS/MS data was used to sequence the cyclic peptides, guided by transcriptomic data for the six novel peptides, and the known sequence for evolidine.

### RNA-seq and transcriptome assembly

Total RNA was isolated from frozen ground leaf tissue of *M. xanthoxyloides* using the mini hot phenol method (51) which in turn is derived from the protocol of Botella *et al*. (52). Total RNA was treated with DNase and subsequently purified with the NucleoSpin RNA Clean-Up kit (Macherey-Nagel). It was validated by both agarose gel electrophoresis and absorbance measurement on a NanoDrop2000 (Thermo Fisher Scientific). RNA-seq libraries were generated using the TruSeq Stranded Total RNA with Ribo-Zero Plant kit according to the manufacturer’s instructions (Illumina) and sequenced on a NextSeq 550 system (Illumina) as single end reads with a length of 75 bp. The dataset contained 9.7 x 10^9^ raw reads and 95% of reads had an average quality score at or above Q30. The raw reads were deposited in the SRA under BioProject accession number PRJNA561449. The leaf transcriptome was assembled using CLC Genomics Workbench 12.0.2 (QIAGEN Aarhus A/S). The raw reads were trimmed to a quality threshold of Q30 and minimum length 15, and assemblies were performed with word sizes 23, 30, 40, 50 and 64 and minimum contig length 200. Other parameters remained at their default values.

### Analysis of data from the Sequence Read Archive

We searched the SRA for RNA-seq and WGS data for other species in the Rutaceae family. Using CLC Genomics Workbench, RNA-seq data from *Aegle marmelos* fruit pulp (SRA run SRR8190707), *Atalantia buxifolia* mature fruit (SRR6349678); *Bergera koenigii* leaf (SRR2970920); *Clausena excavata* leaf (SRR 6438389)*; Zanthoxylum armatum* young leaf (SRR9179743) and *Zanthoxylum bungeanum* young leaf (SRR9179746), and WGS data for *Atalantia buxifolia* (SRR6188460) and *Clausena lansium* (SRR5796634) were downloaded. Data were assembled to a quality threshold Q30 and minimum contig length 100; other parameters were set to their default values, including word size, which the software determined automatically. The assembled transcriptomic and genomic data were subjected to tBLASTn searches using the evolidine and xanthoxycyclin propeptide sequences to find highly similar sequences. The maximum expected value for search results was set to 1 and BLOSUM45 was used as the substitution database.

### Extraction of genomic DNA

Genomic DNA (gDNA) was extracted from 2 g of frozen ground powder of *M. xanthoxyloides* leaves with the DNEasy Plant Mini Kit (QIAGEN) according to the manufacturer’s instructions. Purified DNA was quantified on a NanoDrop 2000 (Thermo Fisher Scientific).

### First strand cDNA synthesis

Complementary DNA (cDNA) was generated from previously extracted total RNA (900 ng) using the SMARTER RACE 5’/3’ kit (Takara Bio USA) according to the manufacturer’s instructions. A poly-T primer was used. The 20 μL reaction was diluted to 110 μL with water.

### PCR and cloning of propeptide genes and transcripts

The DNA template was amplified by the polymerase chain reaction (PCR) using *Pfu* high-fidelity DNA polymerase (Agilent Technologies). Each 50 μL reaction consisted of gDNA (∼27 ng) or cDNA (1 μL) and 1 U *Pfu* DNA polymerase solution, with MgCl2 to a final concentration of 2 mM, 400 μM mixed dNTPs, 10 mM KCl, 10 mM (NH4)2SO4, 20 mM Tris-HCl (pH 8.8), 0.1 mg/mL bovine serum albumin, 0.1% Triton X-100, and forward and reverse primers 0.4 μM (**Table S1**).

PCR amplification was carried out in a Veriti 96-well thermocycler (Applied Biosystems) programmed as follows: 95 °C for 2 min followed by 35 cycles of 95 °C for 30 s; 55 °C for 30 s; 72 °C for 30 s; and finally 72 °C for 10 min. In some cases it was necessary to repeat the PCR a second time using 2 μL of the initial PCR product to produce sufficient DNA for subsequent reactions. PCR products were purified using the QIAquick PCR Purification Kit (QIAGEN) according to the manufacturer’s instructions. DNA was eluted in 30 μL of water after allowing incubation on the column membrane for 1 min and quantified on a NanoDrop 2000. The *Proxanthoxycyclin D* cDNA was Sanger sequenced directly from the PCR product using the same primers as the PCR (**Table S1**).

The other products were cloned as follows: overhanging adenosine bases were added to PCR products in an “A-tailing” reaction using *Taq* DNA polymerase. Each 20 μL reaction consisted of ∼150 ng PCR product, 1 U of *Taq* DNA polymerase, dNTPs to a final concentration of 400 μM, 2 mM MgCl2, 20 mM Tris-HCl (pH 8.8) and 50 mM KCl. Reactions were incubated at 37 °C for 20 min.

Either ∼12.5 ng (*Proxanthoxycyclin C* cDNA), ∼25 ng (*Proxanthoxycyclin B* cDNA) or ∼30 ng (*Proevolidine* and *Proxanthoxycyclin E* gDNA) of the A-tailed PCR product were ligated into 50 ng of the pGEM-T Easy vector (Promega) according to the manufacturer’s instructions. Each reaction was incubated overnight at 4 °C. LB agar plates were prepared containing 100 μg/mL ampicillin and 100 μM isopropyl β-D-1-thiogalactopyranoside (IPTG). Plates were spread with 20 μL of 50 mg/mL 5-bromo-4-chloro-3-indolyl-β-d-galactoside (X-Gal), which was allowed to be absorbed for at least 30 minutes. These plates were inoculated with a culture of TOP10 *E. coli* cells transformed with 2 μL of the ligated vector and incubated overnight at 37 °C. Aliquots of LB (5 mL) were each inoculated with a single colony from a plate and incubated overnight at 37 °C with shaking.

Plasmid DNA was extracted with the GeneJET Plasmid Miniprep Kit (Thermo Fisher Scientific) according to the manufacturer’s instructions. DNA was eluted in 50 μL of water and sent for dideoxy sequencing using the M13F primer of the pGEM-T Easy vector (Australian Genome Research Facility, Perth). The seven evolidine- and proxanthoxycyclin-encoding genes or transcripts from this study were deposited in GenBank under accession numbers MN655991-MN655996 and MT473297.

### NMR Spectroscopy

Xanthoxycyclin samples were prepared by dissolving 2 mg of peptide in 550 μL of H2O/D2O (90:10) at pH ∼3.5. One dimensional ^1^H data, homonuclear ^1^H-^1^H two dimensional TOCSY (37), NOESY (38) and DQF-COSY (Double Quantum Filtered-Correlated Spectroscopy) experiments were recorded at 298 K on a 900 MHz Bruker Avance III spectrometer equipped with a cryoprobe. TOCSY experiments were recorded with 8 scans and 512 increments and two NOESY experiments were recorded with 48 scans and 512 increments with mixing times of 100 ms and 200 ms. Sweep widths of 10 ppm were sufficient to cover all proton resonances. Additionally, a ^1^H-^13^C HSQC spectrum was recorded at natural abundance using 128 scans and 256 increments with a sweep width of 10 ppm in the F2 direct dimension and 80 ppm in the F1 dimension. All data were processed using Topspin 4.0.3 (Bruker), with the solvent signal referenced to 4.77 ppm at 298 K. The spectra were analysed and assigned with the program CARA (Computer Aided Resonance Assignment) (53) using sequential assignment strategies (54). Observed Hα, Cα and Cβ shifts were determined and compared to random coil values (55) for all residues to generate secondary shift graphs. Additionally, amide proton temperature dependence was determined by recording TOCSY data at varying temperatures (288 K, 293 K, 298 K, 303 K, and 308 K). The chemical shift of each HN proton was plotted against temperature to determine the temperature coefficients.

### Solution structure calculations

Integrated peak volumes from NOESY data were used to determine inter-proton distance restraints for xanthoxycyclin D. TALOS-N (39) was used to predict the φ (C^-1^-N-Cα-C) and ψ (N-Cα-C-N^+1^) backbone dihedral angles. Hydrogen bond donors were identified by backbone amide temperature coefficients. Values >-4.6 ppb/K for the coefficient are indicative of hydrogen bond donation (56); hydrogen bond acceptors were identified from preliminary structure calculations (56). Preliminary structures (50) were calculated using torsion angle simulated annealing in CYANA (Combined Assignment and dYnamics Algorithm for NMR Applications) (57). The results of these calculations defined the starting coordinates and distance restraints used for refinement in the program CNS (Crystallography and NMR System) (58). NOE distances, dihedral restraints from TALOS-N and hydrogen bond restraints were all used as input for CNS. Simulated annealing using torsion angle dynamics was performed in CNS to generate 50 structures. All 50 structures were then further minimized in water using Cartesian dynamics to generate the final structures of xanthoxycyclin D. MolProbity (59) was used for stereochemical analysis of the structures via comparison to previously published structures in the PDB. Images of the structure of the xanthoxycyclin D were generated by the program MOLMOL (60). The best 20 structures based on good geometry, no violations above 0.2 Å or 2°, and low energy were chosen to represent the solution structure of xanthoxycyclin D.

The structure of xanthoxycyclin D was submitted to the PDB under the identifier 6WPV and to the Biological Magnetic Resonance Bank (30747).

### Antibacterial and antifungal disc diffusion assays

Synthetic peptides were dissolved in DMSO to a concentration of either 20 mg/mL (evolidine) or 50 mg/mL (xanthoxycyclins D and F).

Antibacterial assays were carried out by a previously described method (8). Briefly, sterile LB agar plates were prepared and evenly inoculated with cultures of *E. coli* K-12 or *B. subtilis* Marburg No. 165 using a sterile swab. The cultures had been prepared in liquid LB medium and diluted to an optical density at 600 nm (OD600) of 0.1. We also prepared sterile 8 mm filter paper discs containing either a positive control (50 μg kanamycin), negative control (5 μL sterile water) or one of the three peptides. Six discs contained 200, 100, 50, 25, 10 or 5 μg of evolidine or 200, 100, 50,.25, 12.5 or 5 μg of xanthoxycyclin D or F. The discs were placed onto the six bacterial plates, incubated overnight at 37 °C and then inspected for any inhibition bacterial growth by the discs.

A similar but simplified disc diffusion assay was carried out against two species of fungi: *Aspergillus fumigatus* and *Candida albicans*. A glycerol stock of each species was streaked onto a sterile yeast extract, peptone and dextrose (YPD) agar plate. Cultures were incubated at 37 °C for ∼48 h before spores were harvested with 3 mL of sterile 0.1% Tween 80. The spore suspensions were then diluted to an OD600 of 0.05 and spread onto fresh YPD plates with a sterile swab. Control discs of sterile 8 mm filter paper were prepared with 50 μg amphotericin B (positive control) or 5 μL DMSO (negative control). Another three discs contained 200 μg of each peptide. A set of discs was placed onto each of the two fungal plates and incubated for ∼24 h at 37 °C before inspection for any inhibition of fungal growth.

## Author Contributions

J.S.M. conceived the study and purified RNA; D.C. provided plant material; T.C. performed RNA and peptide extractions; O.B. and J.W. performed RNA-seq; M.F.F. and T.C. assembled the transcriptomic and genomic data, analyzed the assemblies and performed PCR; M.F.F. performed LC-MS, analyzed LC-MS data and analyzed dideoxy sequencing data; C.P. synthesized peptide; C.P. and K.J.R. performed NMR studies; M.F.F., C.P., K.J.R. and J.S.M. wrote the manuscript with help from all other authors.

## Acknowledgments

The authors thank Dr Fanie Venter of James Cook University for collecting the plant material used in this study. The authors thank Dr Nicolas L. Taylor of the School of Molecular Sciences at the University of Western Australia for providing advice and assistance in mass spectrometry aspects of this project. The authors acknowledge the facilities, and the scientific and technical assistance of the Australian Microscopy & Microanalysis Research Facility at the Centre for Microscopy, Characterisation & Analysis, The University of Western Australia, a facility funded by the University, State and Commonwealth governments.

## Supplementary Information

### Supplementary Tables

**Table S1.**
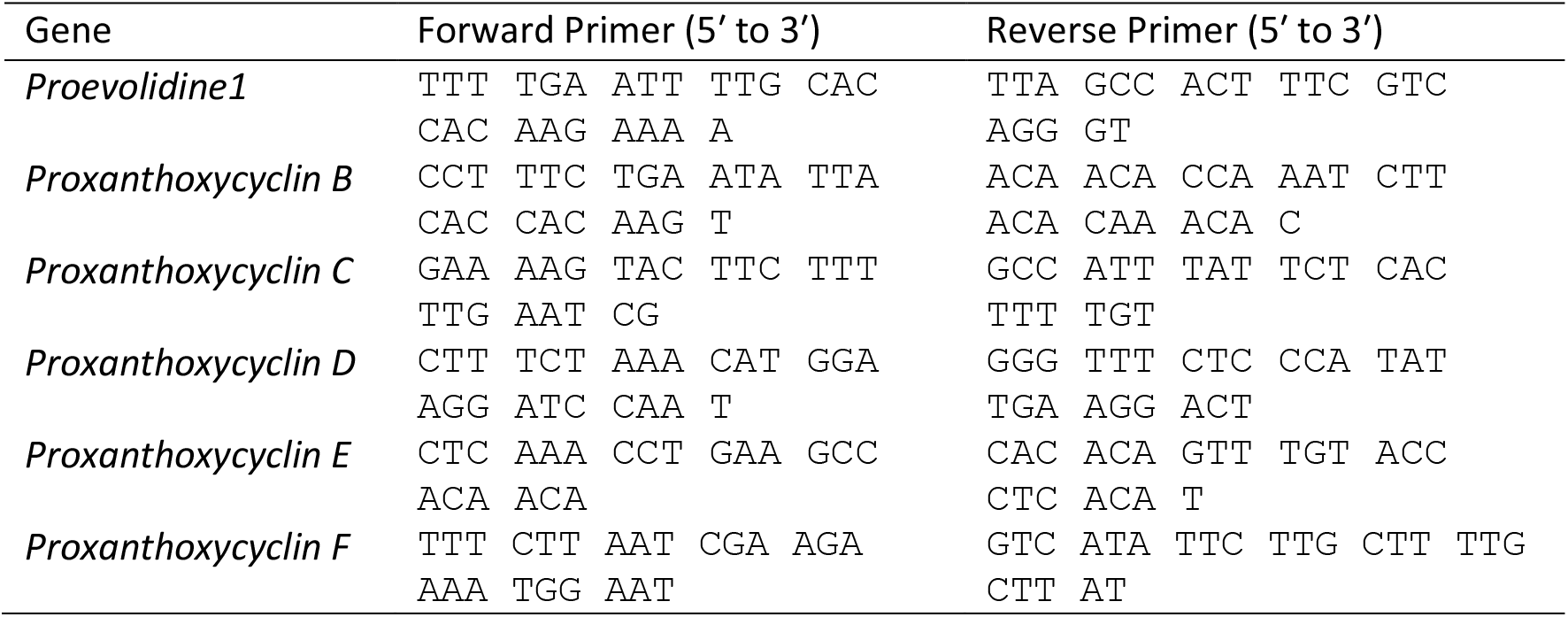
Primers used for PCR amplification of genes encoding orbitides in *M. xanthoxyloides*.

### Supplementary Figures

**Figure S1.**
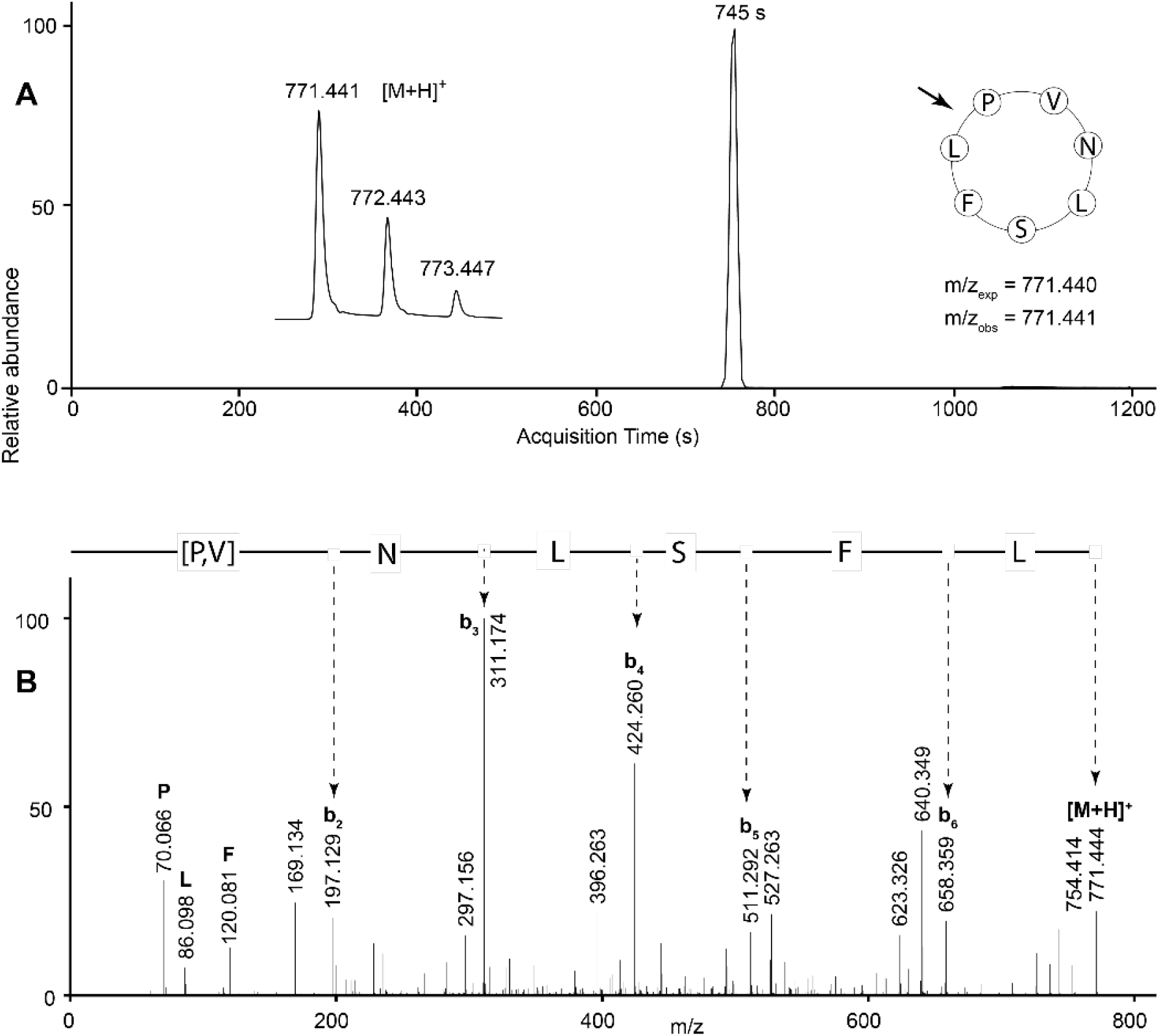
LC-MS/MS confirmation of evolidine. (A) Extracted ion chromatogram at *m/z* 771.440. Inset left: mass spectrum at the chromatogram peak. Inset right: cyclic peptide sequence with MS/MS ring cleavage point shown by an arrow, expected *m/z* (*m/z*_exp_) and observed *m/z* (*m/z*_obs_). (B) MS/MS spectrum of protonated molecule at *m/z* 771.440. A single strong sequence of b ions is derived from initial ring opening between Leu and Pro residues.

**Figure S2.**
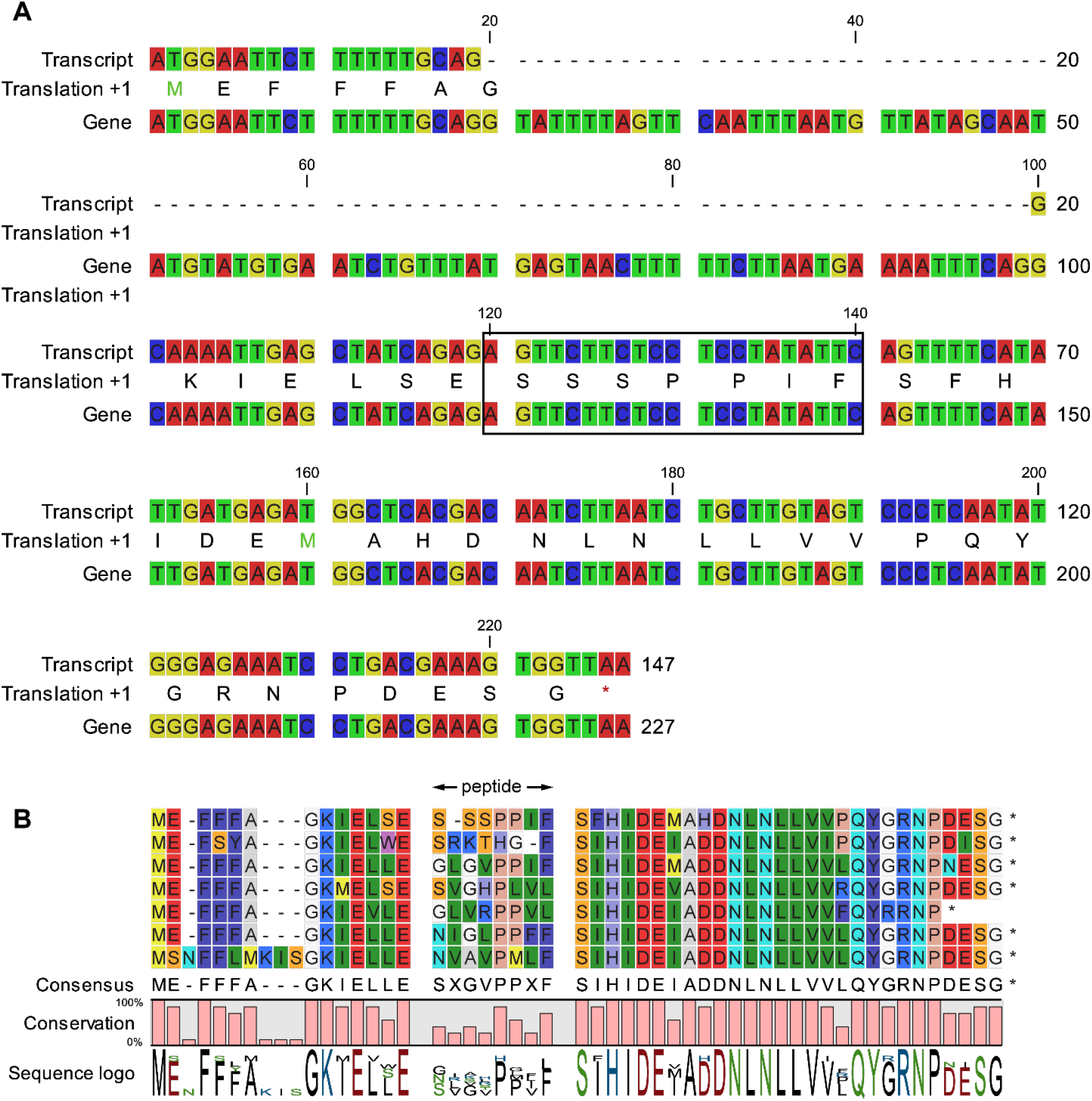
Alignment of the *proxanthoxycyclin E* transcript with genetic data. (A) The Sanger-sequenced gene data matches perfectly with the RNA-seq transcript, except that the gene contains an 80 bp intron. The translated peptide sequence is shown between the nucleotide sequences; xanthoxycyclin E is shown within a box. (B) Alignment of the *xanthoxycyclin E* translated gene sequence (top line) with other potential gene sequences amplified by PCR from genomic DNA template using the same primer pair. All contain an 80 bp intron.

**Figure S3.**
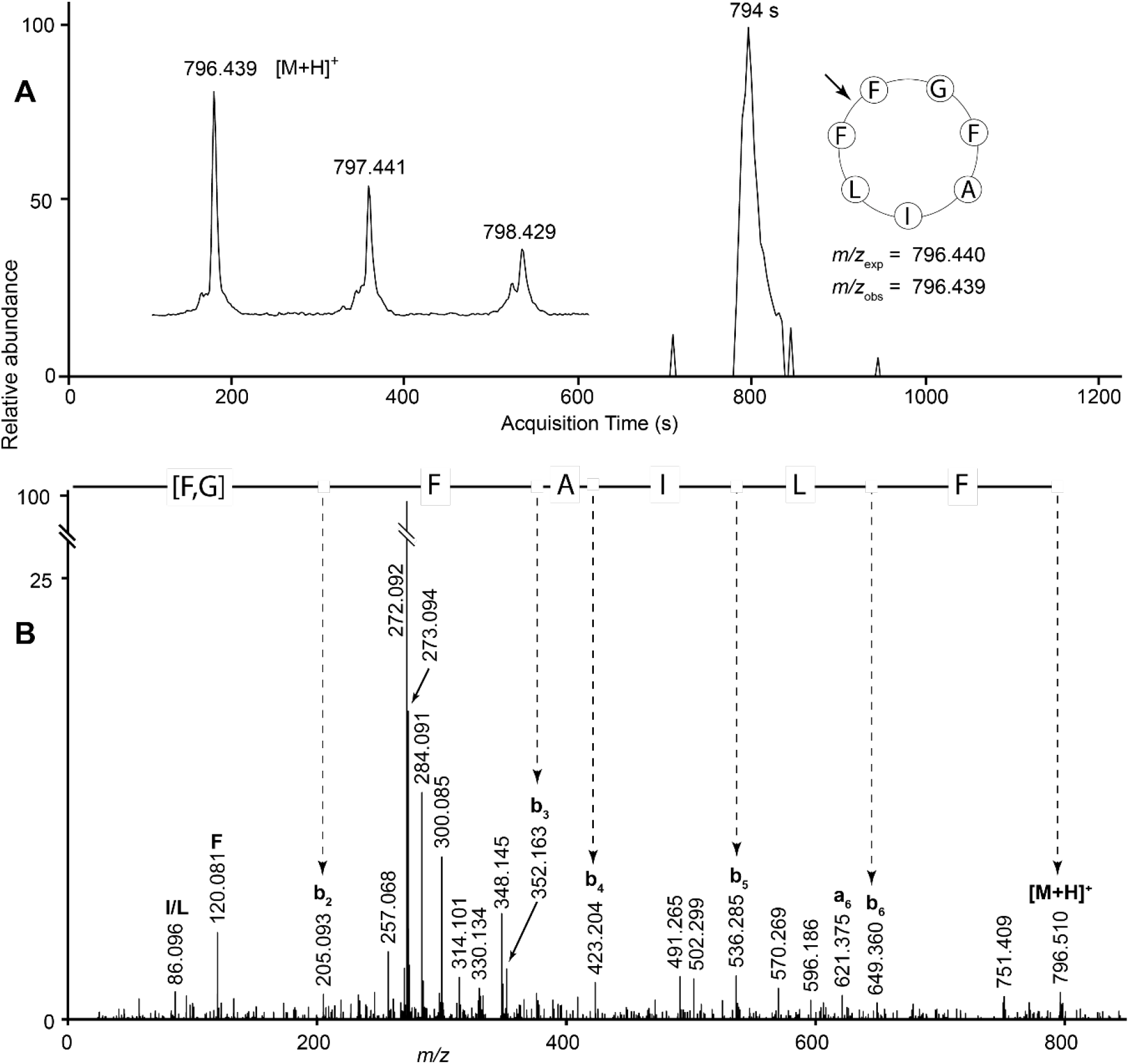
LC-MS/MS data for xanthoxycyclin B. (A) Extracted ion chromatogram at *m/z* 796.440. Inset left: mass spectrum at the chromatogram peak. Inset right: cyclic peptide sequence with MS/MS ring cleavage point shown by an arrow, expected *m/z* (*m/z*_exp_) and observed *m/z* (*m/z*_obs_). (B) MS/MS spectrum of protonated molecule at *m/z* 796.440. A strong sequence of b ions is derived from initial ring opening between two Phe residues.

**Figure S4.**
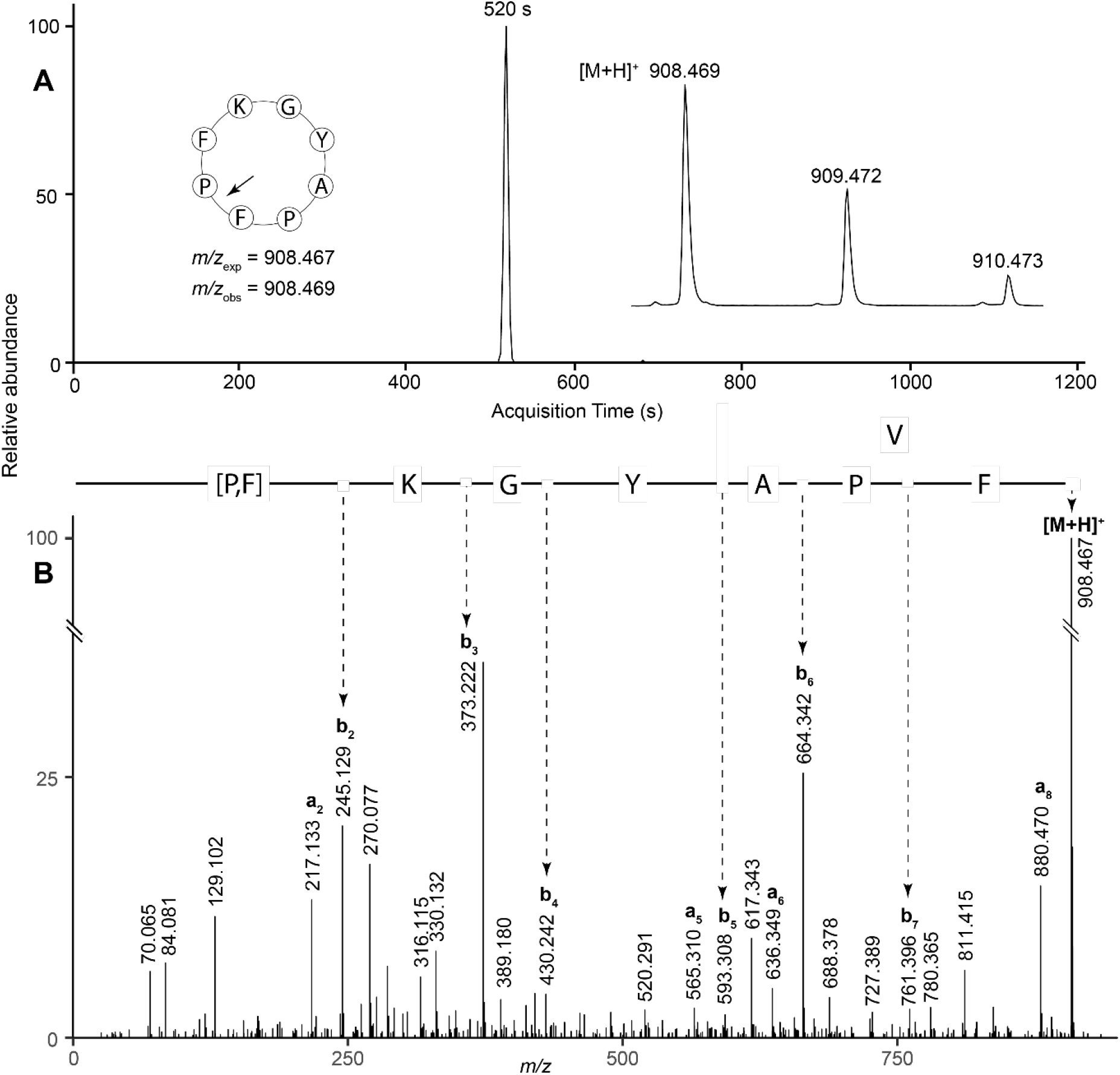
LC-MS/MS data for xanthoxycyclin C. (A) Extracted ion chromatogram at *m/z* 908.467. Inset left: cyclic peptide sequence with MS/MS ring cleavage point shown by an arrow, expected *m/z* (*m/z*_exp_) and observed *m/z* (*m/z*_obs_). Inset right: mass spectrum at the chromatogram peak. (B) MS/MS spectrum of protonated molecule at *m/z* 908.467. A strong sequence of b ions is derived from initial ring opening between Phe and Pro residues.

**Figure S5.**
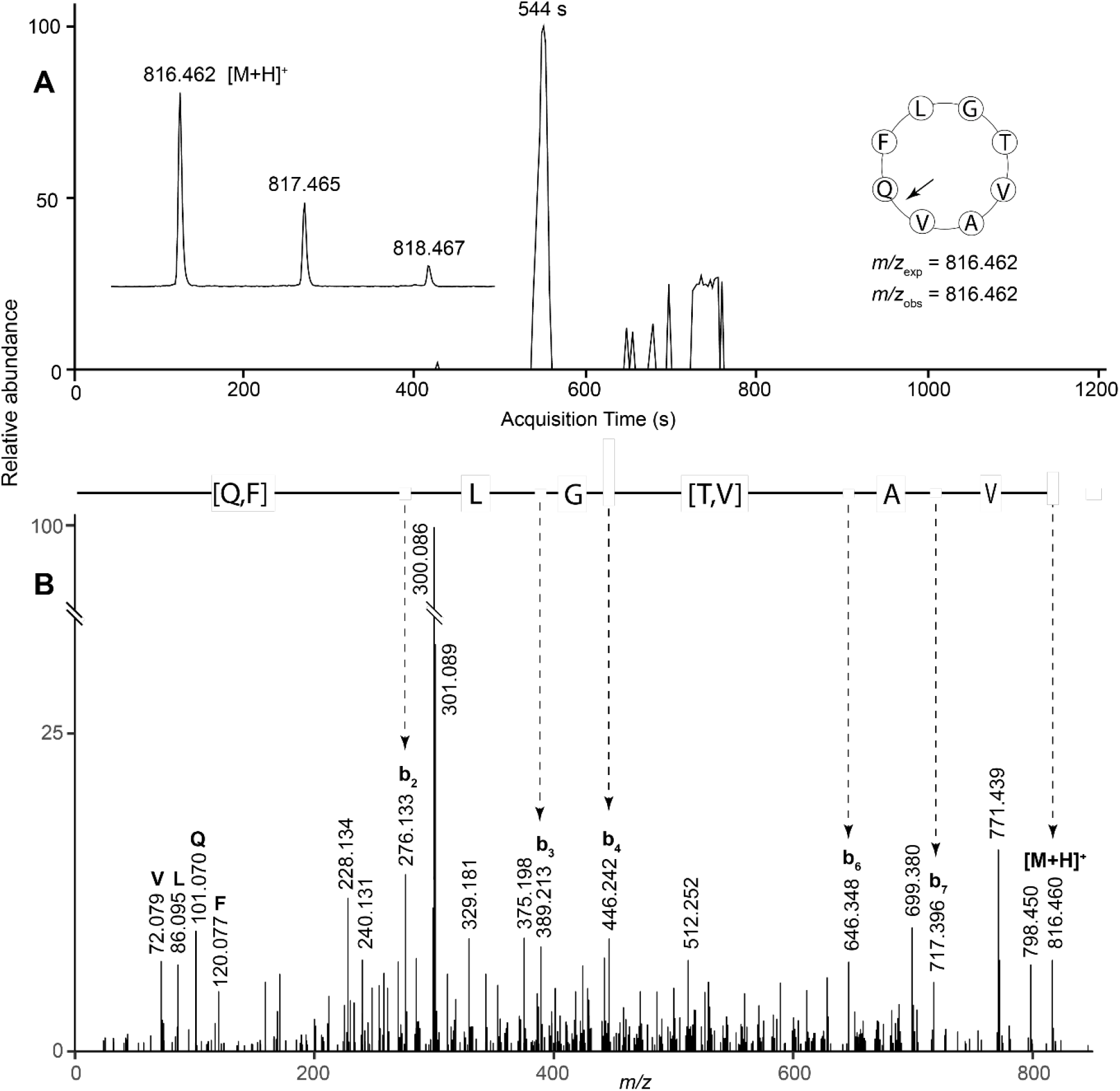
LC-MS/MS data for xanthoxycyclin D. (A) Extracted ion chromatogram at *m/z* 816.462. Inset left: mass spectrum at the chromatogram peak. Inset right: cyclic peptide sequence with MS/MS ring cleavage point shown by an arrow, expected *m/z* (*m/z*_exp_) and observed *m/z* (*m/z*_obs_). (B) MS/MS spectrum of protonated molecule at *m/z* 816.462. A sequence of b ions is derived from initial ring opening between the Val and Gln residues.

**Figure S6.**
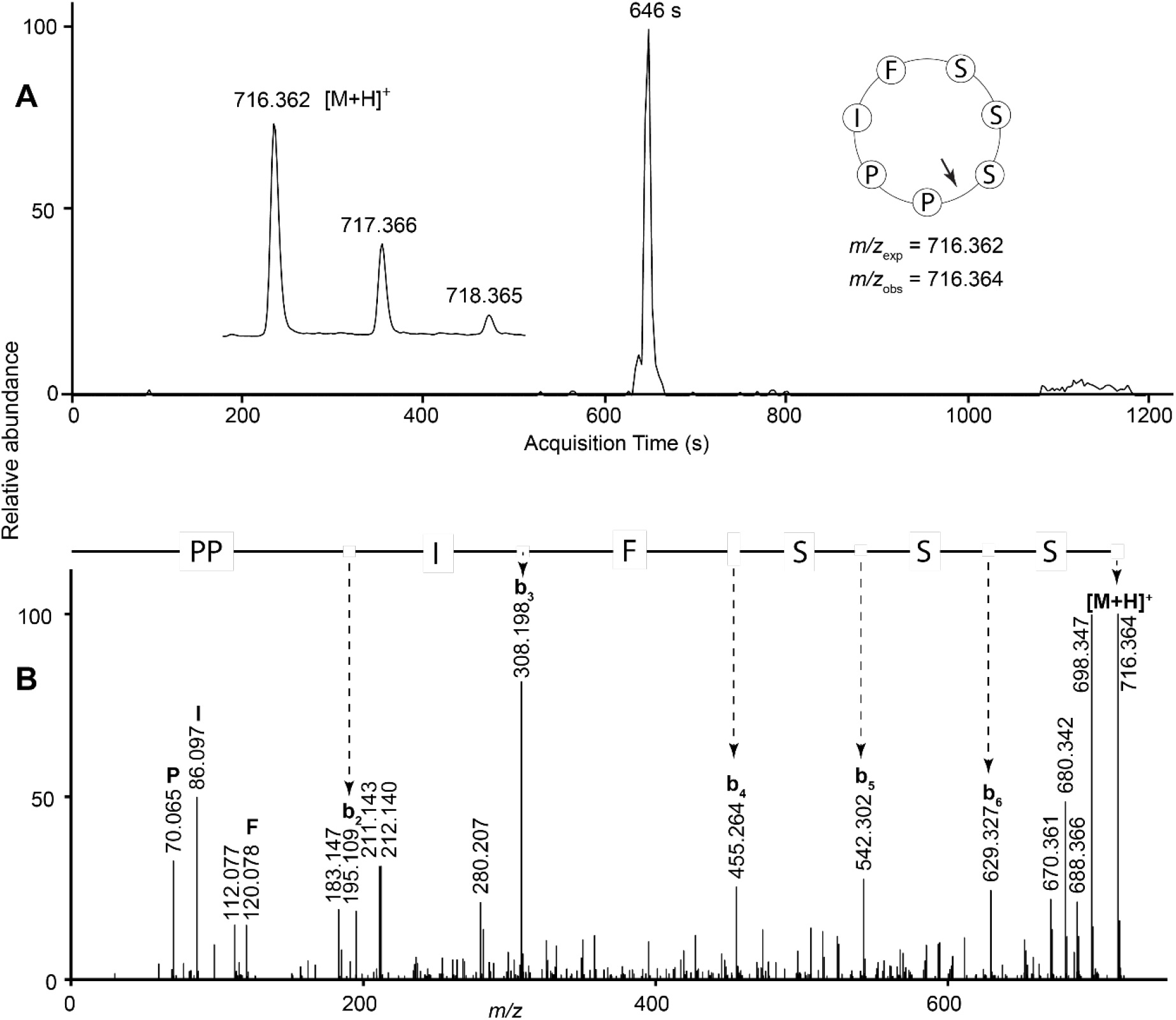
LC-MS/MS data for xanthoxycyclin E. (A) Extracted ion chromatogram at *m/z* 716.362. Inset left: mass spectrum at the chromatogram peak. Inset right: cyclic peptide sequence with MS/MS ring cleavage point shown by an arrow, expected *m/z* (*m/z*_exp_) and observed *m/z* (*m/z*_obs_). (B) MS/MS spectrum of protonated molecule at *m/z* 716.362. A strong sequence of b ions is derived from initial ring opening between Ser and Pro residues.

**Figure S7.**
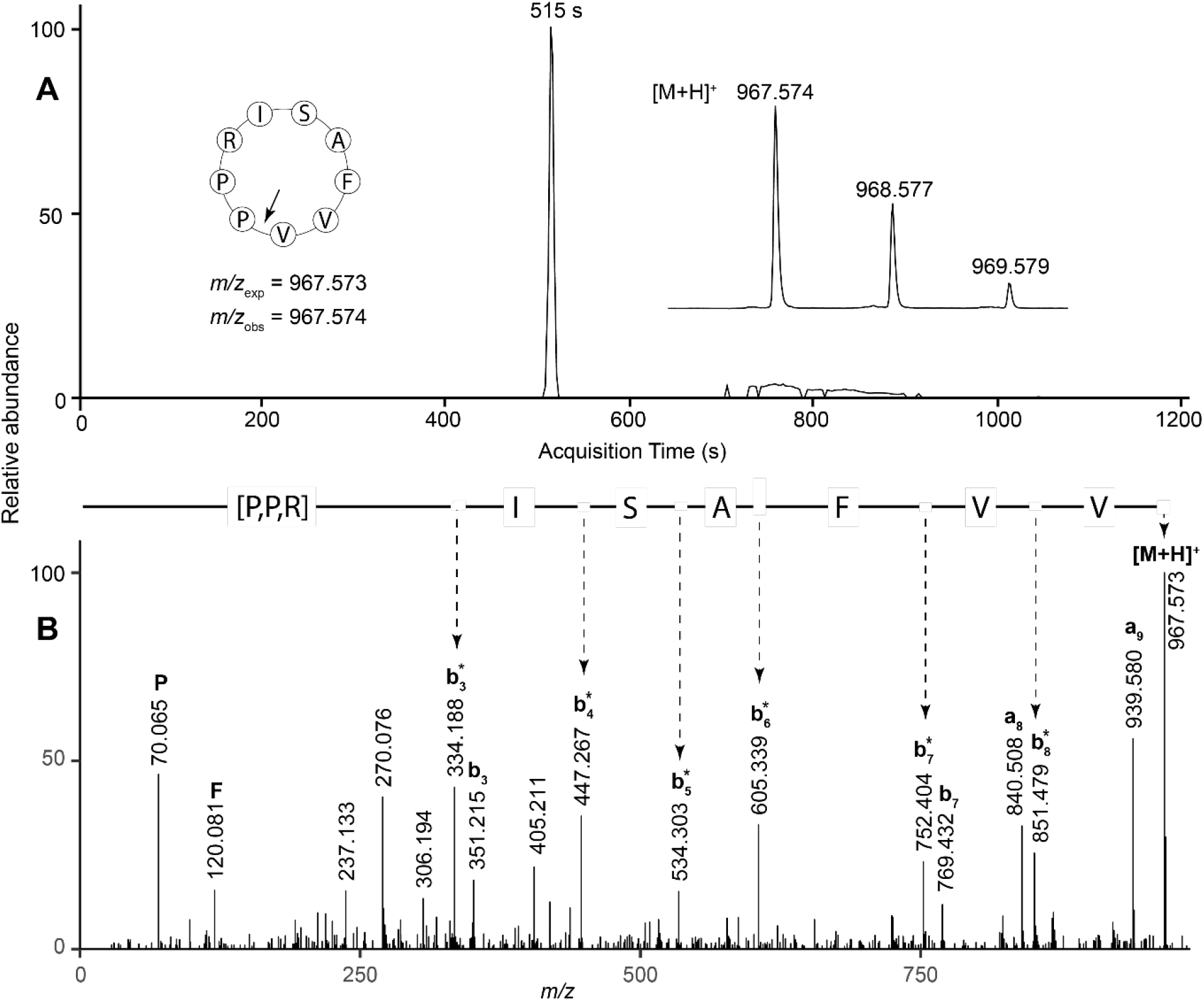
LC-MS/MS data for xanthoxycyclin F. (A) Extracted ion chromatogram at *m/z* 967.573. Inset left: cyclic peptide sequence with MS/MS ring cleavage point shown by an arrow, expected *m/z* (*m/z*_exp_) and observed *m/z* (*m/z*_obs_). Inset right: mass spectrum at the chromatogram peak. (B) MS/MS spectrum of protonated molecule at *m/z* 967.573. A strong sequence of b* ions is derived from initial ring opening between Val and Pro residues. b* ions are b ions with the neutral loss of NH3 from the Arg residue.

**Figure S8.**
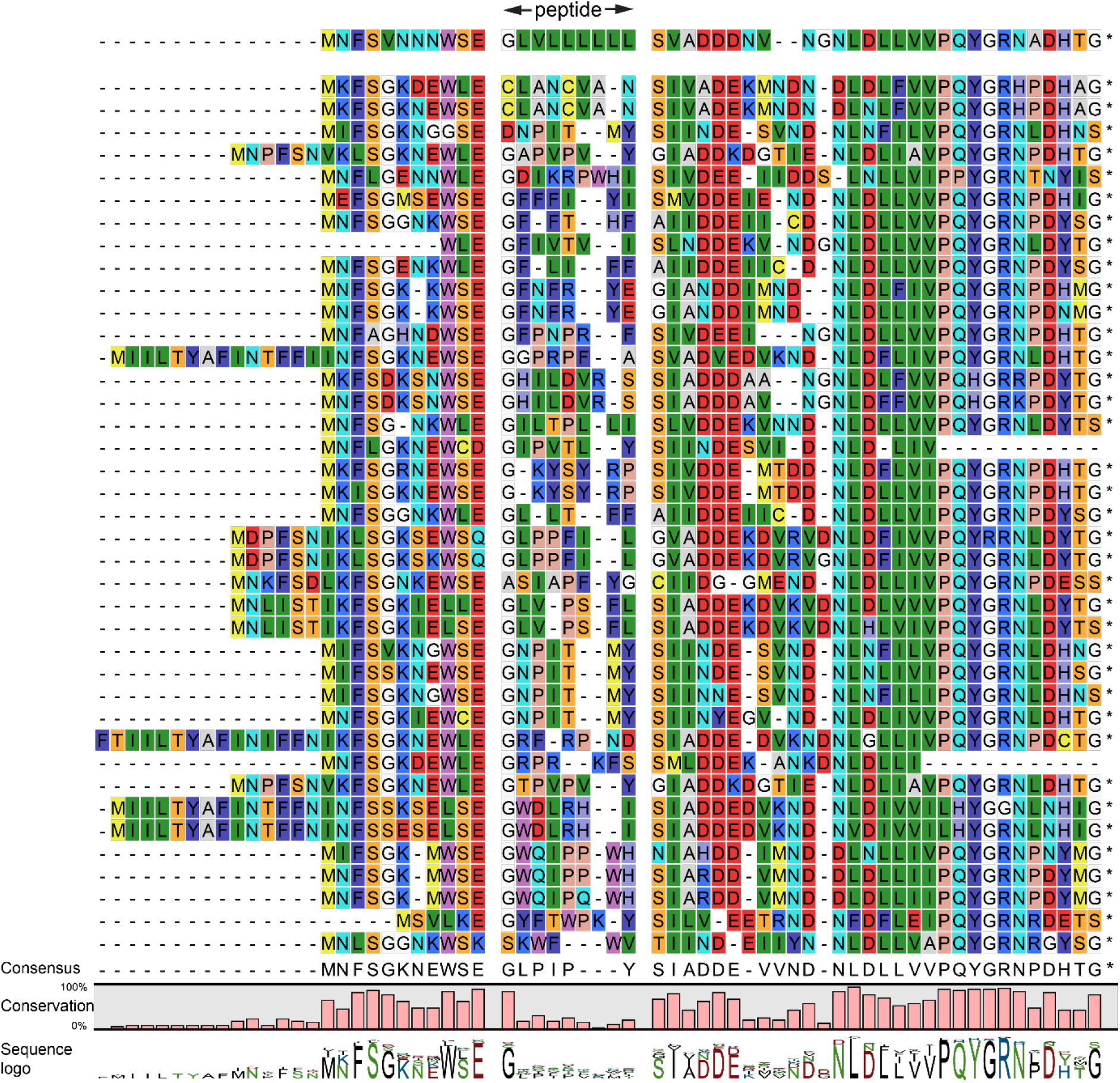
Alignments of translated coding sequences for putative *Atalantia buxifolia* propeptides. Sequences are shown with background RasMol colours, and the putative peptide is marked and spaced from the surrounding sequence. Asterisks represent stop codons. Below the alignment are the consensus sequence, degree of conservation and a sequence logo. The first sequence appears to encode clausenlanin B, a peptide previously discovered in another Rutaceae species, *Clausena lansium* (32).

**Figure S9.**
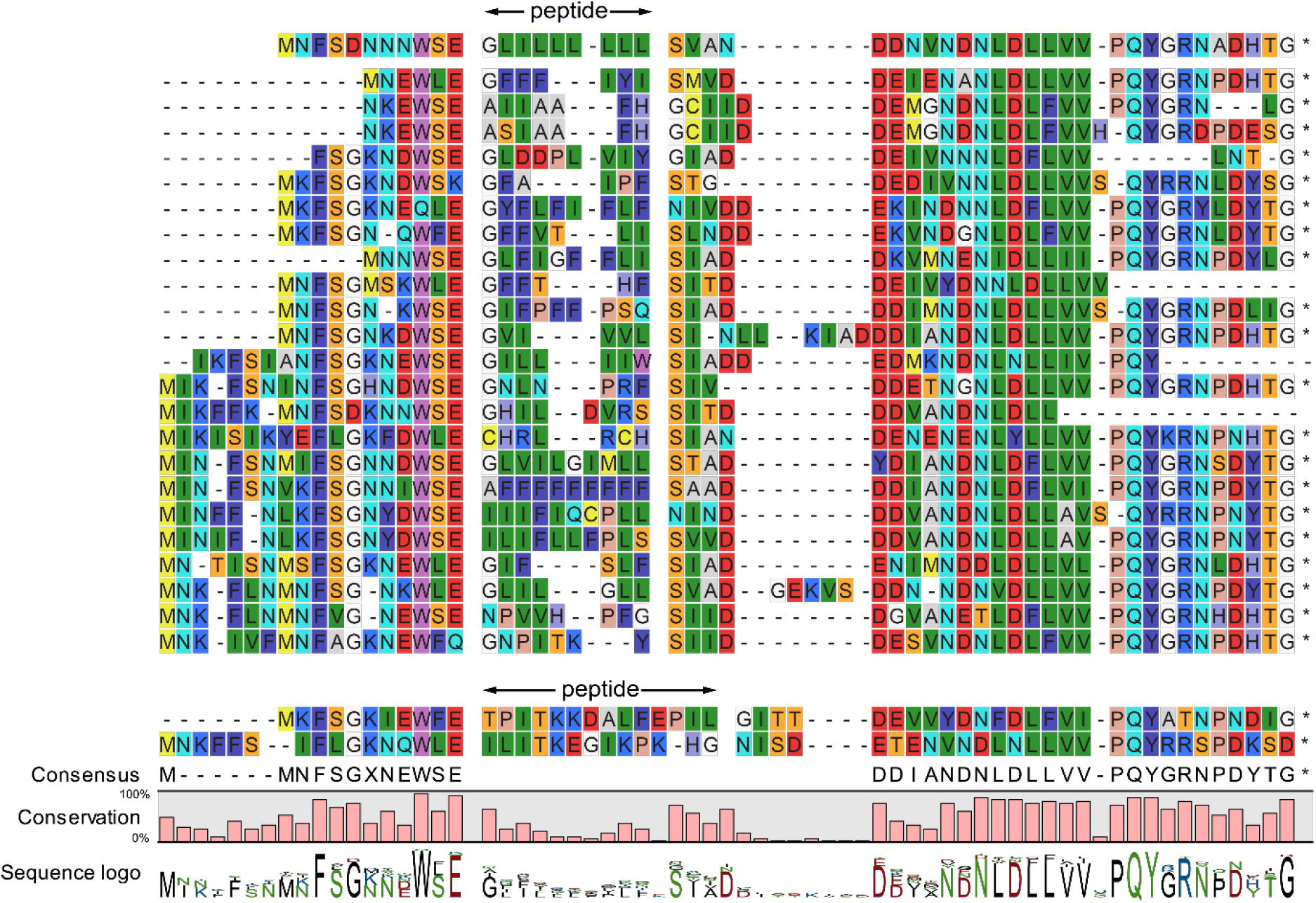
Alignments of translated coding sequences for putative *Clausena lansium* propeptides. Sequences are shown with background RasMol colours, and the putative peptide is marked and spaced from the surrounding sequence. Asterisks represent stop codons. Below the alignment are the consensus sequence, degree of conservation and a sequence logo. The first sequence appears to encode the known peptide clausenlanin A (32).

**Figure S10.**
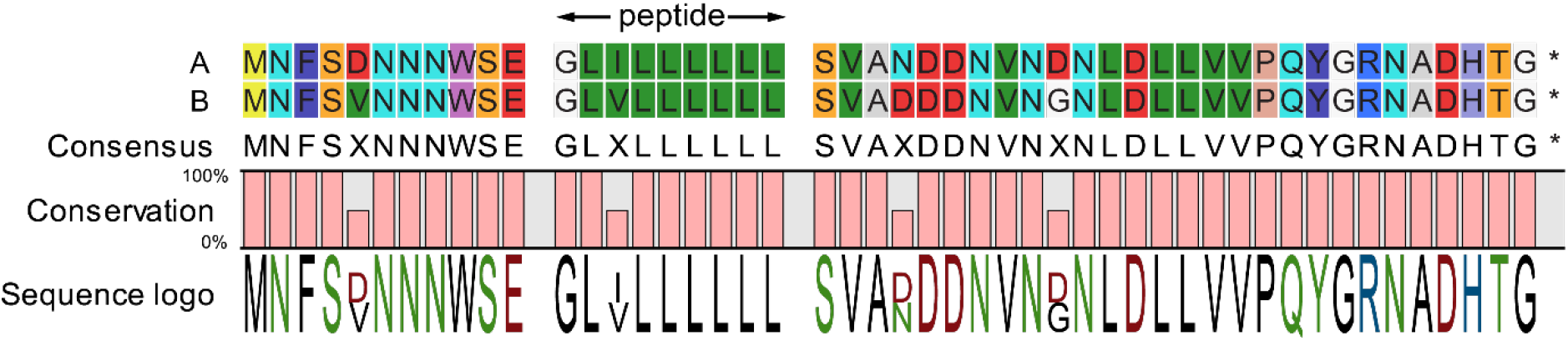
Alignments of translated potential coding sequences for clausenlanins A and B. Note the high degree of conservation between sequence A, found in genomic data from *Clausena lansium*, and sequence B, found in *Atalantia buxifolia*. Sequences are shown with background RasMol colours, and the putative peptide is marked and spaced from the surrounding sequence. Below the alignment are the consensus sequence, degree of conservation and a sequence logo.

**Figure S11.**
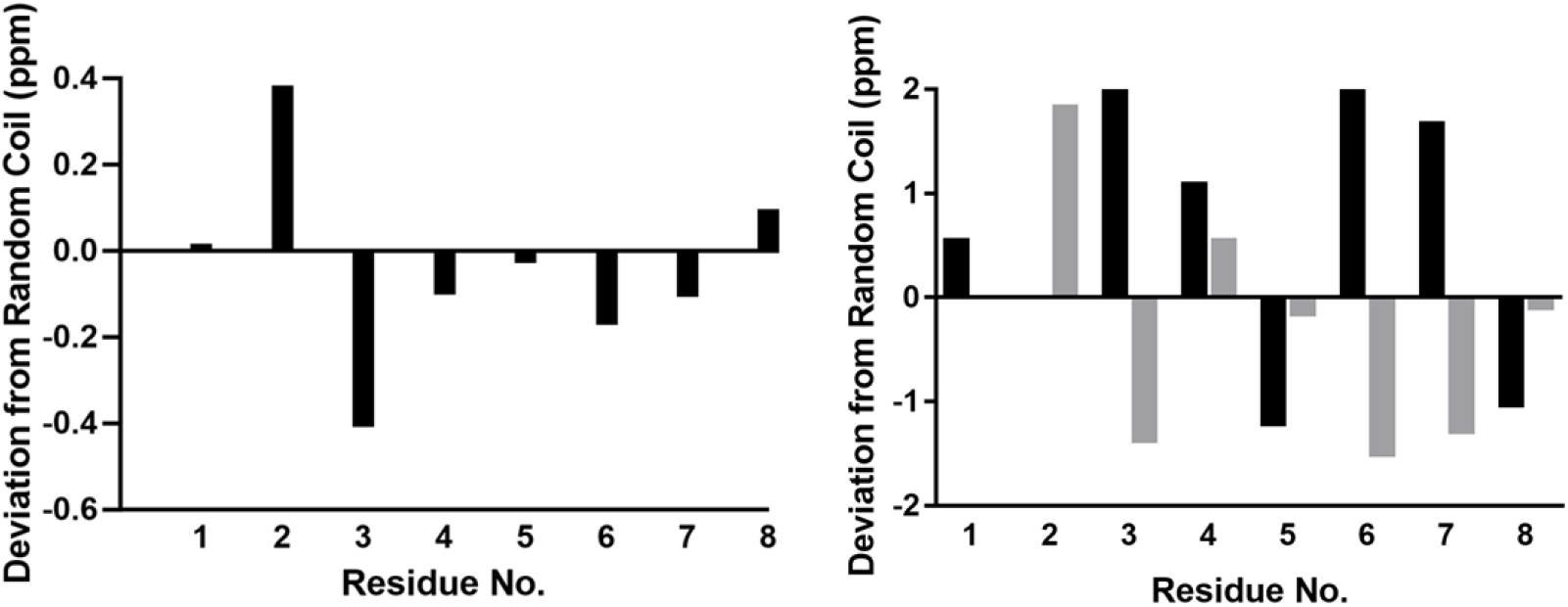
Secondary ^1^Hα, ^13^Cα and ^13^Cβ shifts of xanthoxycyclin D. Observed deviations from random coil suggest an ordered backbone region. Left graph displays the secondary ^1^Hα shifts in black. Right graph displays the secondary ^13^Cα shifts in black and ^13^Cβ shifts in grey.

**Figure S12.**
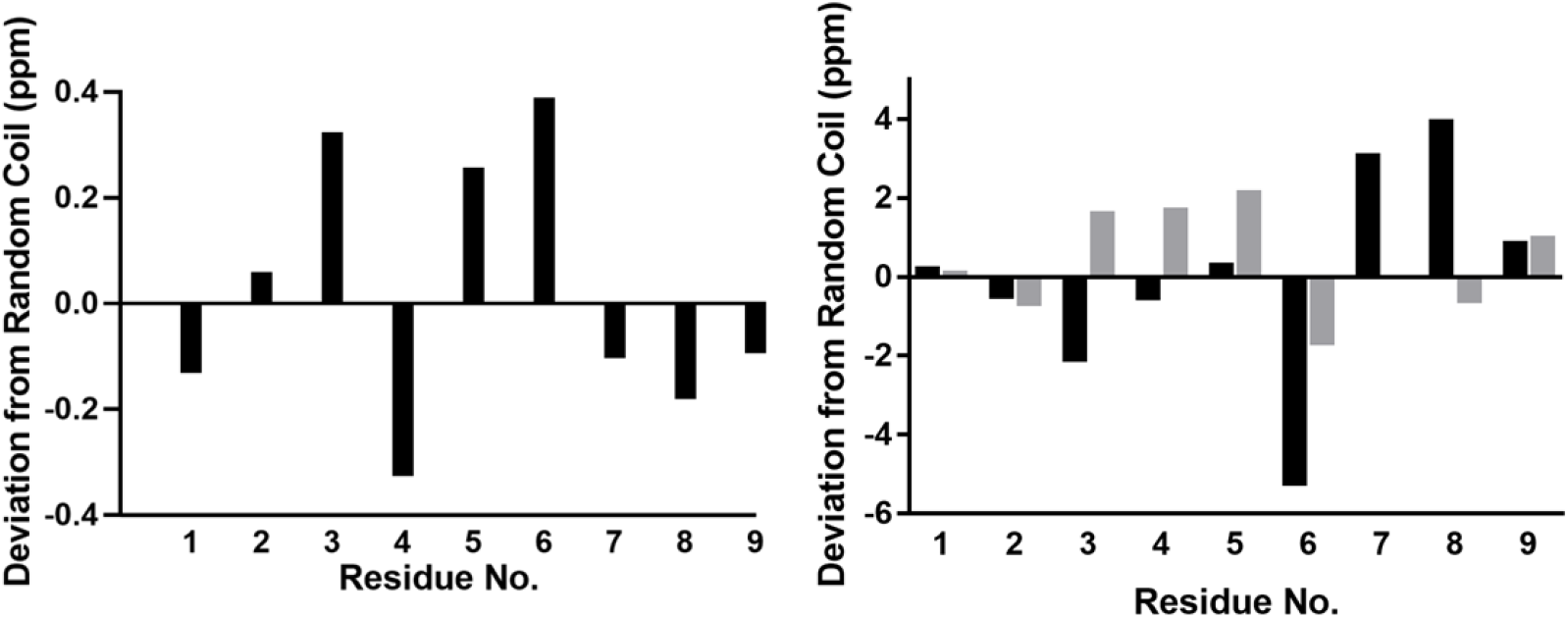
Secondary ^1^Hα, ^13^Cα and ^13^Cβ shifts of xanthoxycyclin F. Observed deviations from random coil suggest an ordered backbone region. Left graph displays the secondary ^1^Hα shifts in black. Right graph displays the secondary ^13^Cα shifts in black and ^13^Cβ shifts in grey.

**Figure S13.**
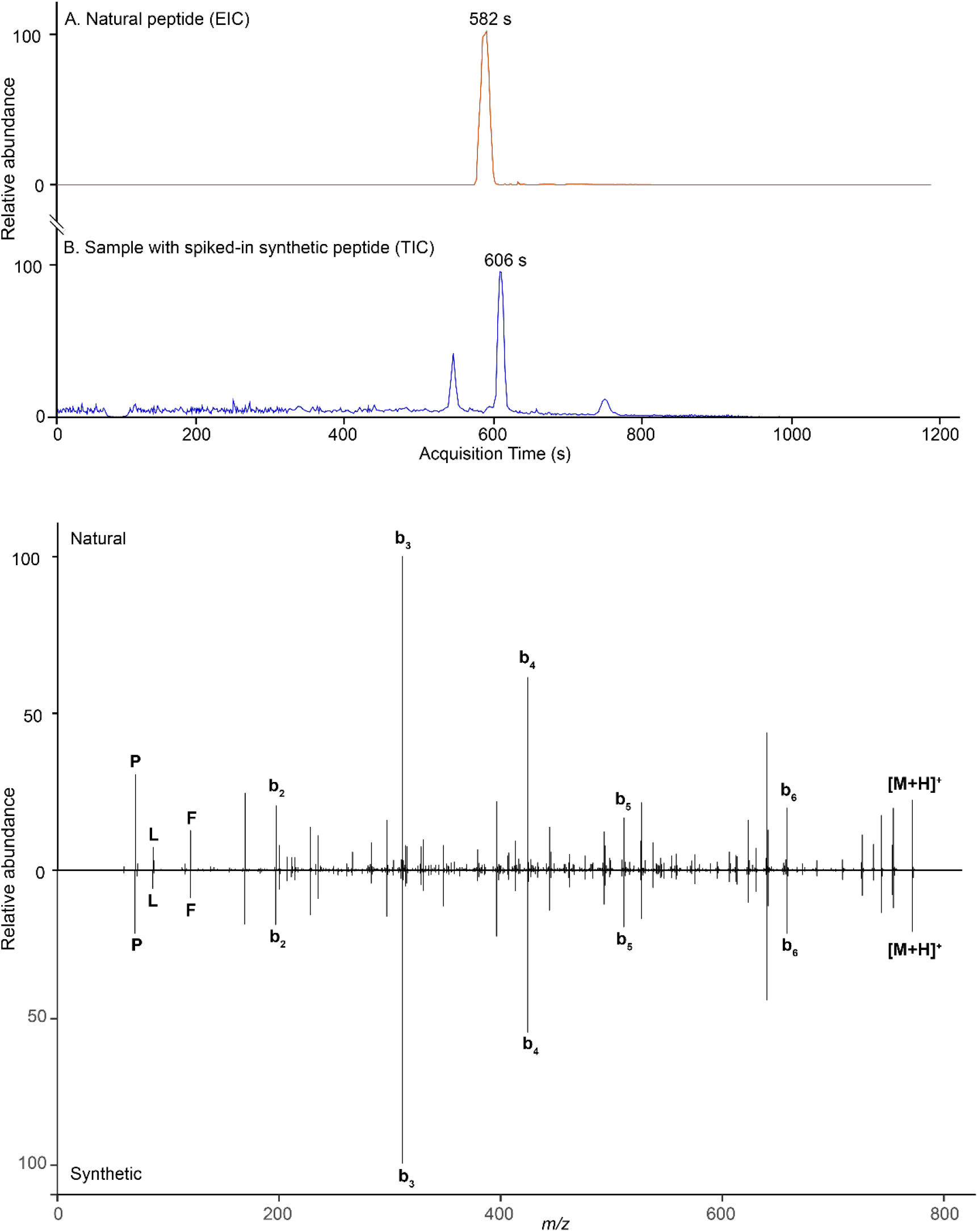
LC-MS/MS comparison of natural and synthetic evolidine. (A) Comparison of the extracted ion chromatogram (EIC) of evolidine in the native peptide extract with the total ion current chromatogram (TIC) of the native extract with synthetic evolidine spiked-in. The acquisition (retention) times for evolidine are similar. (B) A mirror plot comparing the MS/MS spectra of natural evolidine (top) and the synthetic version (bottom). The spectra are labelled with the protonated molecule ([M+H]^+^), the b ion series and immonium ions (marked with their single letter amino acid code).

**Figure S14.**
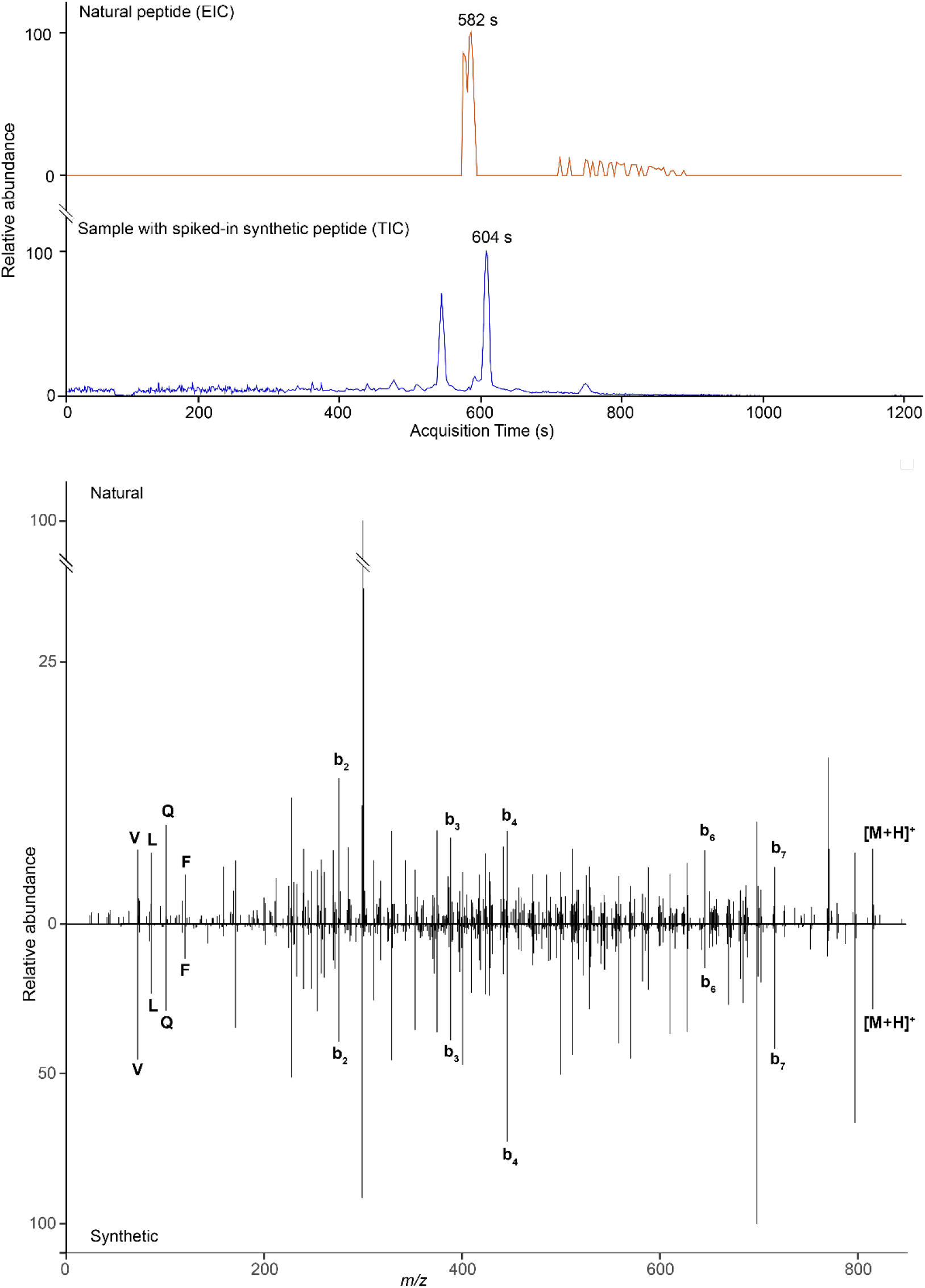
LC-MS/MS comparison of natural and synthetic xanthoxycyclin D (cyclo-GTVAVQFL). (A) Comparison of the EIC of xanthoxycyclin D in the native peptide extract with the TIC of the native extract with synthetic xanthoxycyclin D spiked-in. The acquisition (retention) times for xanthoxycyclin D are similar. (B) A mirror plot comparing the MS/MS spectra of natural xanthoxycyclin D (top) and the synthetic version (bottom). The spectra are labelled with the protonated molecule ([M+H]^+^), the b ion series and immonium ions (marked with their single letter amino acid code).

**Figure S15.**
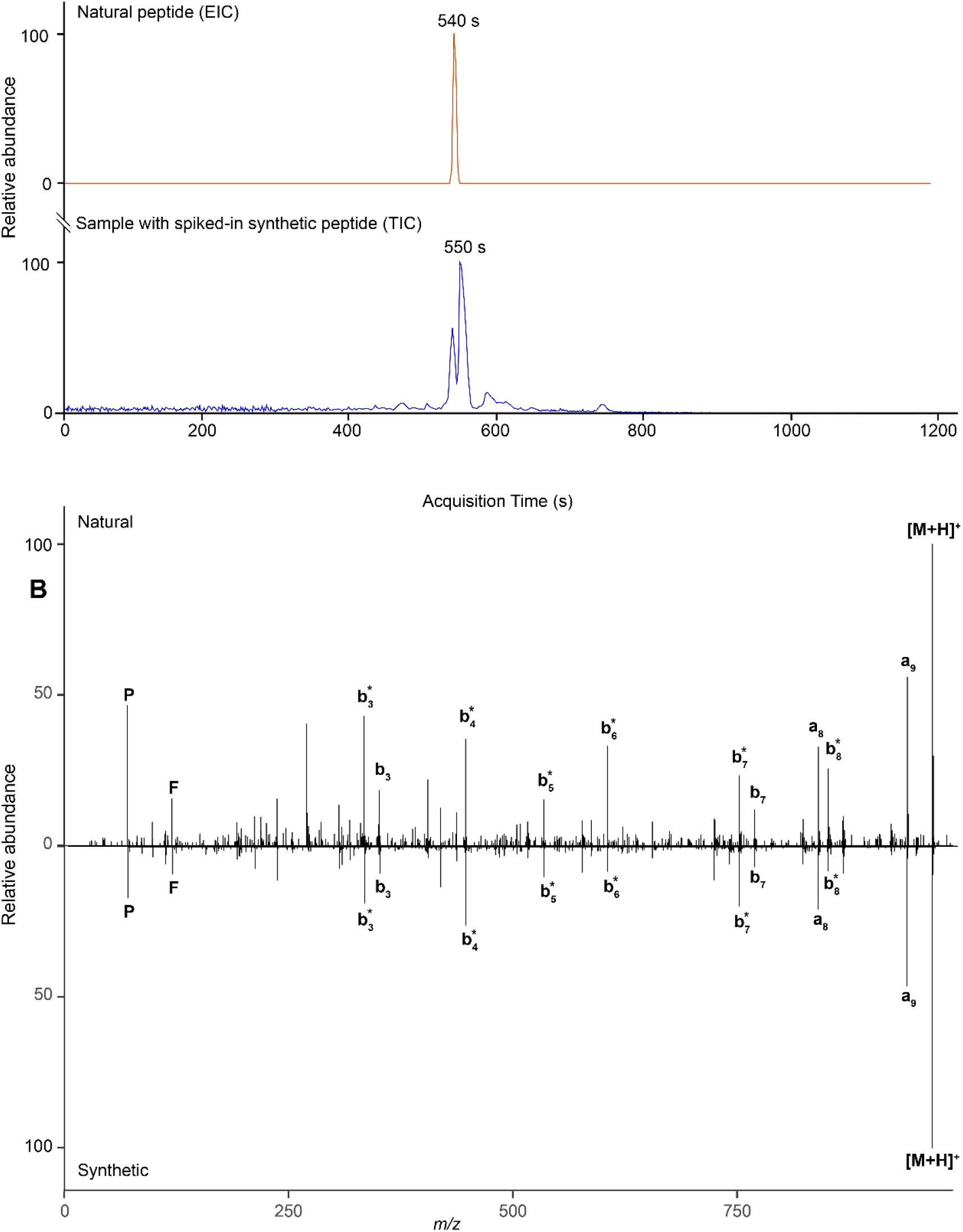
LC-MS/MS comparison of natural and synthetic xanthoxycyclin F (cyclo-AFVVPPRIS). (A) Comparison of the EIC of xanthoxycyclin F in the native peptide extract with the TIC of the native extract with synthetic xanthoxycyclin F spiked-in. The acquisition (retention) times for xanthoxycyclin F are similar. (B) A mirror plot comparing the MS/MS spectra of natural xanthoxycyclin F (top) and the synthetic version (bottom). The spectra are labelled with the protonated molecule ([M+H]^+^), the a, b and b* ion series and immonium ions (marked with their single letter amino acid code).

**Figure S16.**
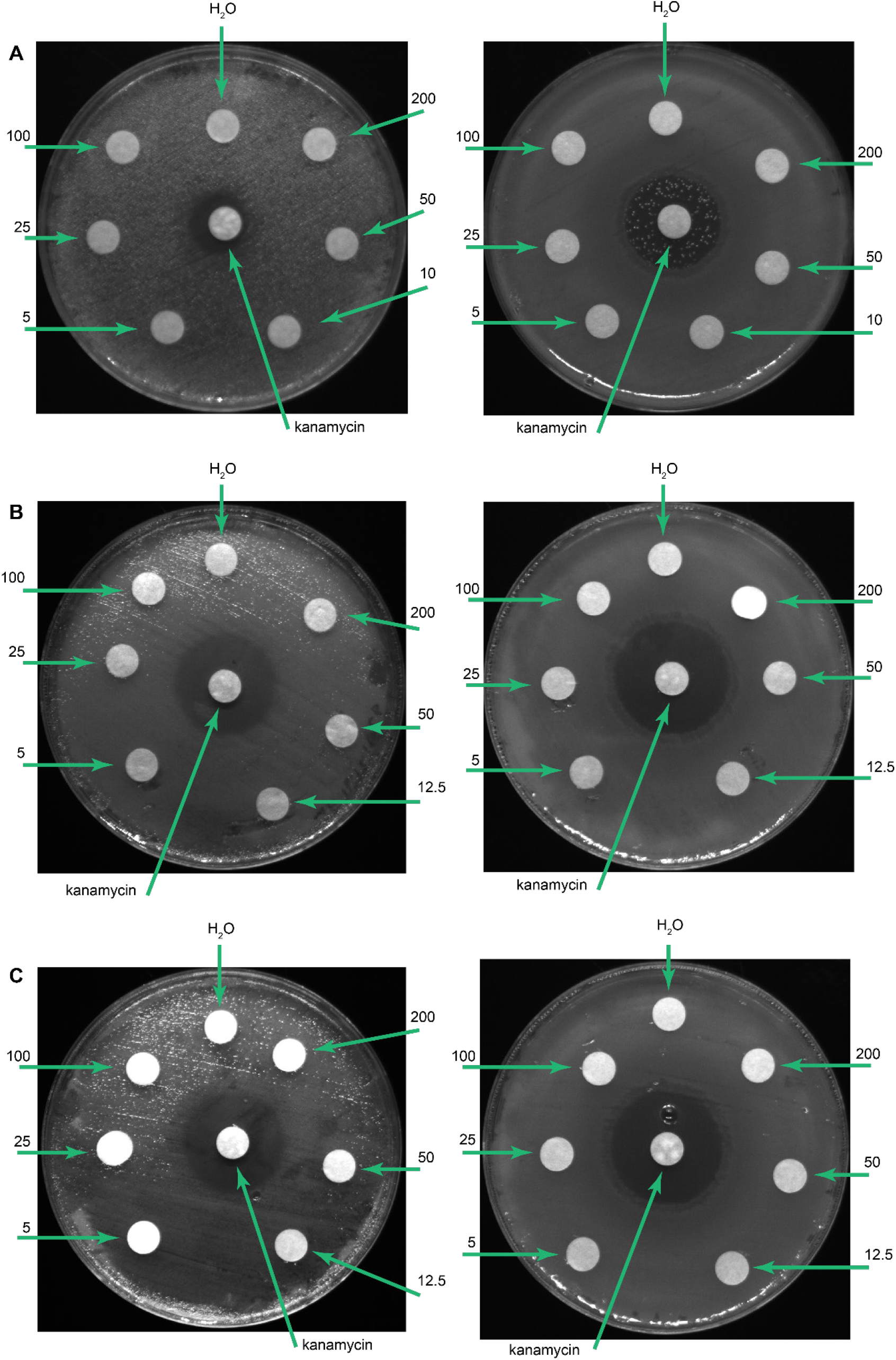
Results of disc diffusion assays against bacteria. Left: *E. coli* K12; Right: *B. subtilis* Marburg No. 168 (A) evolidine, (B) xanthoxycyclin D, (C) xanthoxycyclin F. The numbers represent the amount of peptide applied to the disc (μg). Positive control is kanamycin (50 μg) and negative control is DMSO (5 μL). None of the three peptides inhibited bacterial growth.

**Figure S17.**
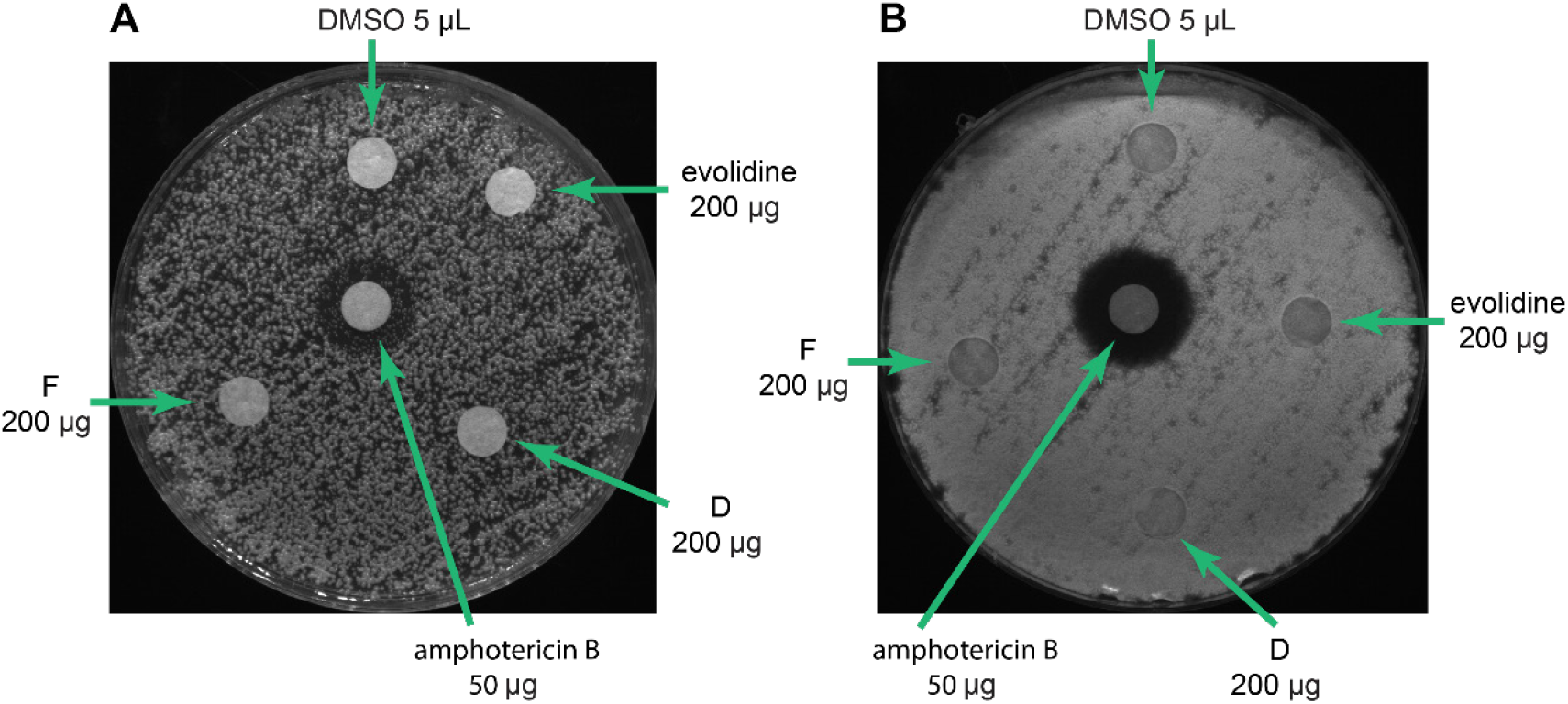
Results of disc diffusion assays against fungi. (A) *C. albicans;* (B) *A. fumigatus*. Positive control is amphotericin B (50 μg) and negative control is DMSO (5 μL). D and F are xanthoxycyclin D and xanthoxycyclin F. None of the three peptides inhibited fungal growth.

**Figure S18.**
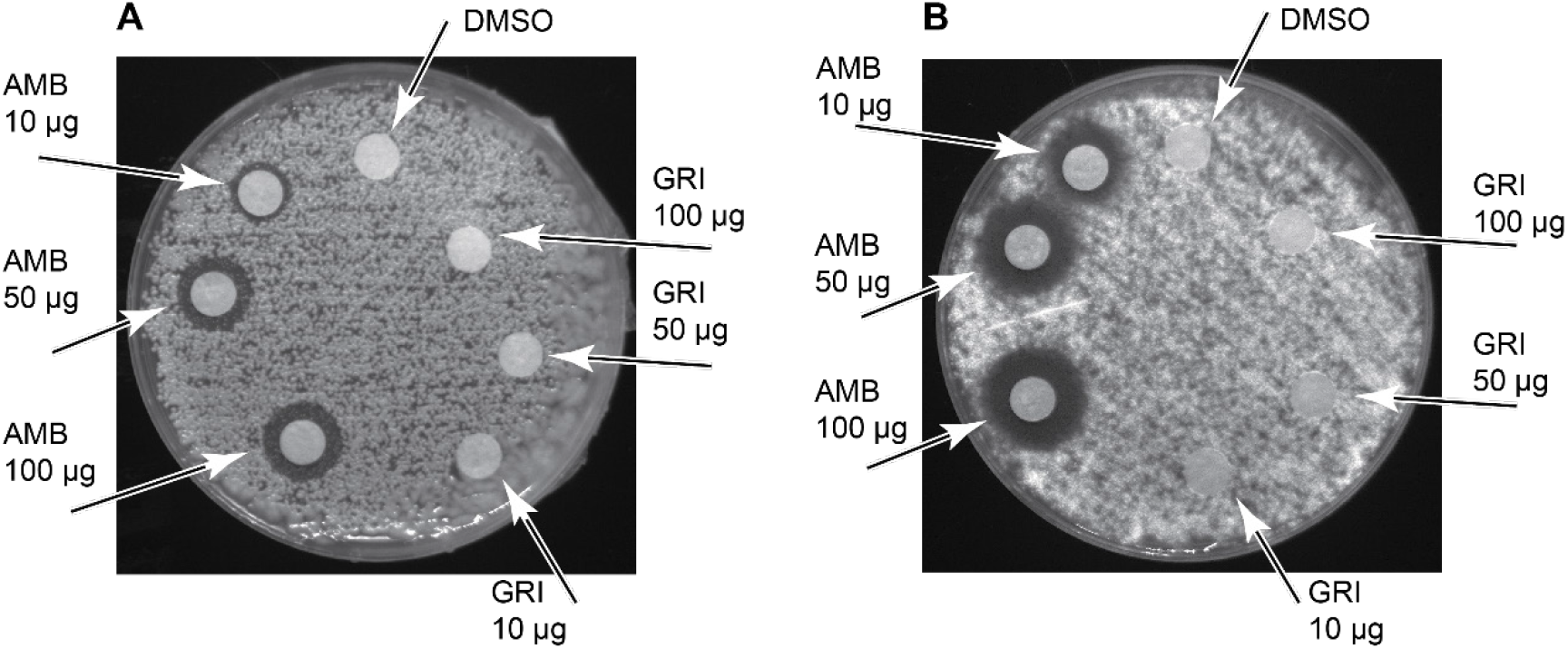
Selection of an antifungal agent as a positive control for antifungal assays. (A) *C. albicans;* (B) *A. fumigatus*. AMB is amphotericin B and GRI is griseofulvin. DMSO (5 μL) was used as a negative control.

